# Chromosome-level assemblies of multiple Arabidopsis genomes reveal hotspots of rearrangements with altered evolutionary dynamics

**DOI:** 10.1101/738880

**Authors:** Wen-Biao Jiao, Korbinian Schneeberger

## Abstract

We report chromosome-level, reference-quality assemblies of seven *Arabidopsis thaliana* accessions selected across the global range of this predominately ruderal plant. Each genome revealed between 13-17 Mb rearranged and 5-6 Mb novel sequence introducing copy-number changes in ∼5,000 genes, including ∼1,900 genes which are not part of the current reference annotation. Analyzing the collinearity between the genomes revealed ∼350 regions (4.1% of the euchromatin) where accession-specific tandem duplications destroyed the syntenic gene order between the genomes. These *hotspots of rearrangements* were characterized by the loss of meiotic recombination in hybrids within these regions and the enrichment of genes implicated in biotic stress response. Together this suggests that hotspots of rearrangements are governed by altered evolutionary dynamics as compared to the rest of the genome, which are based on new mutations and not on the recombination of existing variation, and thereby enable a quick response to the ever-evolving challenges of biotic stress.

---

The individual genomes of sexually reproducing species are typically highly collinear to enable physical exchange of alleles (chromosome arms) during meiosis. This exchange ensures the generation of diversity and the removal of deleterious alleles^1–3^ and at the time protects the offspring from major mutations changing the karyotype of a genome^4,5^. Despite the obvious importance to preserve a common karyotype, the presence of genomic rearrangements suggests that the genomes are in fact not entirely collinear. Genomic rearrangements (and the resulting lack of allelic exchange) have been shown to contribute to population diversification including the evolution of different sexes^6,7^ or life-history traits^8^.

But even though the absence of collinearity can have drastic effects, there is hardly anything known about the actual degree of collinearity within populations as most of the current genome studies are not based on chromosome-level assemblies. The first complete assembly of a plant genome was the reference sequence of *A. thaliana* (Col-0), which was based on a minimal tiling path of BACs sequenced with Sanger technology^9^. Since then multiple hundred Arabidopsis genomes have been studied, however, most of these studies relied on short-read based resequencing or reference-guided assembly, where the identification of genomic rearrangements remained challenging^10–16^. In contrast, reference-independent, chromosome-level assemblies with almost complete reconstruction of the nucleotide sequence enable accurate identification of all sequence differences and would therefore reveal the degree of synteny across the genome^17^. So-far, however, there are only few whole-genome *de-novo* assemblies for *A. thaliana* available and the assemblies have not been thoroughly compared to each other^18–21^.

Using deep PacBio (45-71x) and Illumina (56-78x) whole-genome shotgun sequencing, we assembled the genomes of seven accessions from geographically diverse regions including An-1 (Antwerpen, Belgium), C24 (Coimbra, Portugal), Cvi-0 (Cape Verde Islands), Eri-1 (Eringsboda, Sweden), Kyo (Kyoto, Japan), L*er* (Gorzów Wielkopolski, Poland) and Sha (Shahdara, Tadjikistan) (Supplementary Table S1) (see Methods). The assembly of L*er* was already described in the context of the development of a whole-genome comparison tool used in this study^17^. The seven accessions (together with the reference accession Col-0) were initially used as the founder lines of Arabidopsis Multi-parent Recombination Inbreeding Lines (AMPRIL)^22^ population and were selected to maximize the genetic diversity in this set. The contig assemblies featured N50 values from 4.8 - 11.2 Mb and chromosome-normalized L50 (CL50)^23^ values of 1 or 2 indicating that nearly all chromosome arms were assembled into a few contigs only (Fig. 1, Table 1 and Supplementary Table 2). We arranged 43 - 73 contigs per genome to chromosome-level assemblies based on homology to the reference sequence. Even though these assemblies do rely on the reference sequence, we would like to point out that the sequence assembly itself was independent of the reference sequence, and that the contigs were large in general, implying that it is unlikely that we misplaced any of the contigs. To confirm this, we compared two of the chromosome-level assemblies with three different genetic maps, where we did not find even a single mis-placed contig (Supplementary Table 3). The seven chromosome-level assemblies reached a total length of 117.7 - 118.8 Mb, which is very similar to the 119.1 Mb of the reference sequence (Table 1) and even included parts of the highly complex regions of centromeres, telomeres and rDNA clusters (Supplementary Table 4 and 5). The remaining unanchored contigs had a total length of 1.5 - 3.3 Mb and consisted almost entirely of repeats, which agrees with gaps between the contigs, which were mostly introduced due to repetitive regions (Supplementary Table 6). Overall, we annotated 27,098 to 27,574 protein-coding genes in each of the assemblies, which is similar to the 27,445 genes annotated in the reference sequence^24^ (Table 1 and Supplementary Table 7-9) (see Methods).

**Table 1.**
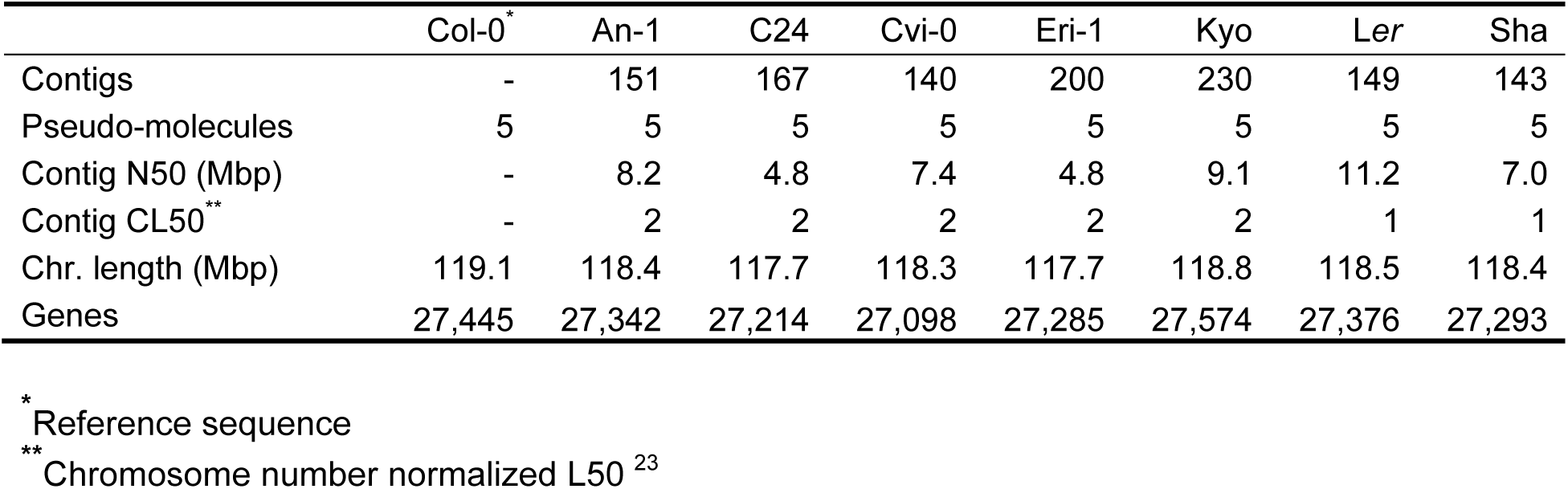
Genome assembly and annotation of seven *A. thaliana* accessions.

|  | Col-0<sup>*</sup> | An-1 | C24 | Cvi-0 | Eri-1 | Kyo | Ler | Sha |
| --- | --- | --- | --- | --- | --- | --- | --- | --- |
| Contigs | - | 151 | 167 | 140 | 200 | 230 | 149 | 143 |
| Pseudo-molecules | 5 | 5 | 5 | 5 | 5 | 5 | 5 | 5 |
| Contig N50 (Mbp) | - | 8.2 | 4.8 | 7.4 | 4.8 | 9.1 | 11.2 | 7.0 |
| Contig CL50<sup>**</sup> | - | 2 | 2 | 2 | 2 | 2 | 1 | 1 |
| Chr. length (Mbp) | 119.1 | 118.4 | 117.7 | 118.3 | 117.7 | 118.8 | 118.5 | 118.4 |
| Genes | 27,445 | 27,342 | 27,214 | 27,098 | 27,285 | 27,574 | 27,376 | 27,293 |

<sup>\*</sup> Reference sequence
<sup>\*\*</sup> Chromosome number normalized L50<sup>23</sup>

**Fig. 1.**
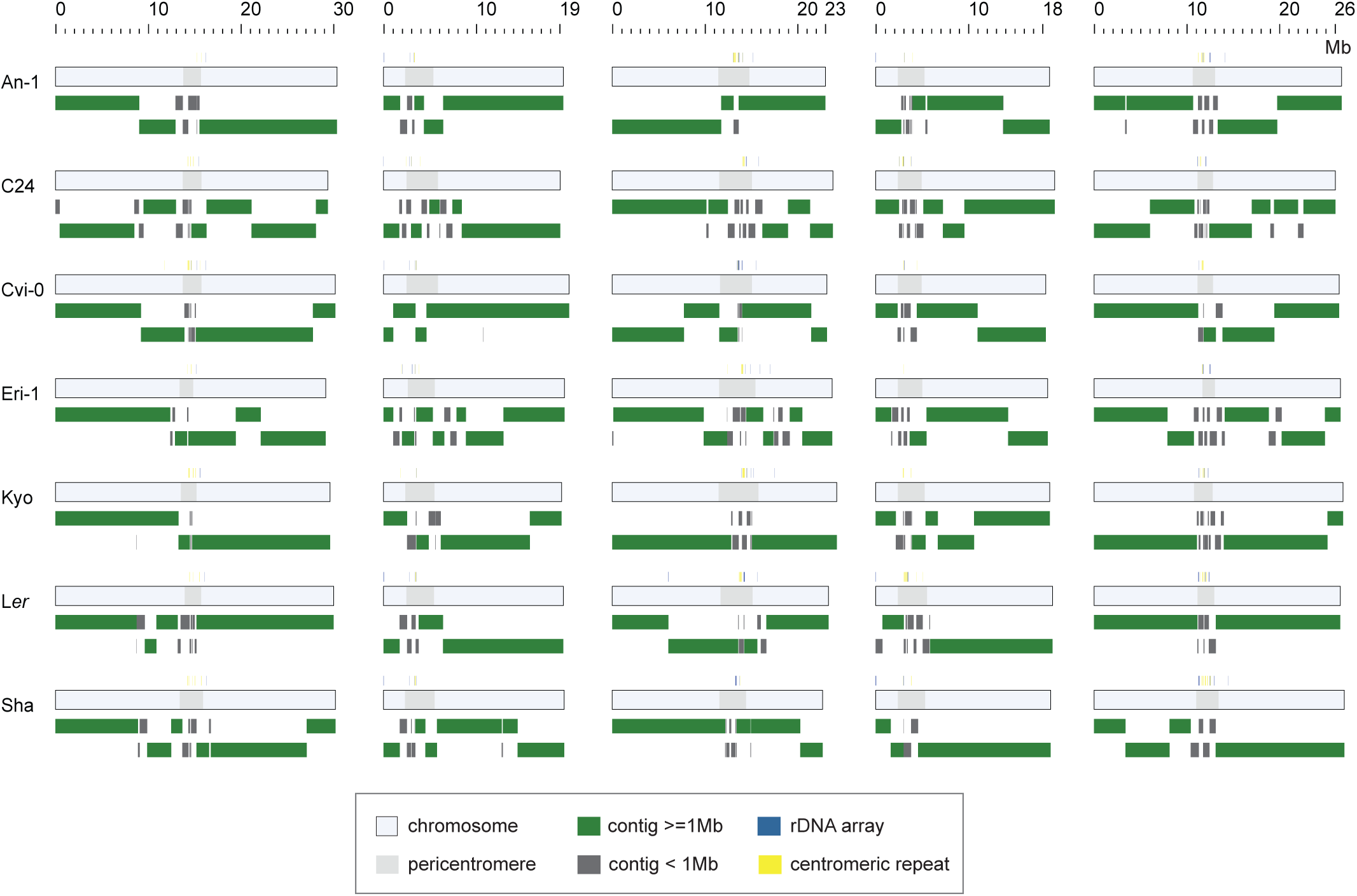
Chromosome-level genome assemblies of seven *A. thaliana* accessions. The light grey bars outline each of the chromosomes, whereas the dark gray inlays show the extend of each of the pericentromeric regions. The contig arrangements of the chromosome assemblies is shown in green for contigs > 1 Mb and dark grey for contigs < 1 Mb. The location of centromeric tandem repeat arrays and rDNA clusters within the assemblies are marked by yellow and blue boxes above each of the chromosomes.

By comparing each of the new assemblies against the reference sequence using the whole-genome comparison tool *SyRI* (V1.1)^17^, we found 102.2 - 106.6 Mb of collinear regions and 12.6 - 17.0 Mb of rearranged regions in each of the genomes (Fig. 2a). The rearrangements included 1.5 - 4.2 Mb (33 - 46) inversions, 1.8 – 2.9 Mb (729 - 1,192) translocations, and - most abundant - polymorphic duplications, which comprised 7.2 – 8.7 Mb (4,288 – 5,150) within each of the individual genomes (Supplementary Table 10). Similar to small-scale sequence variation^25^, rearrangements were not evenly distributed along the chromosomes, but were enriched in pericentromeres (Supplementary Table 11). Their lengths ranged from a few dozen bp to hundreds of kb and even Mb scale (Fig. 2b), including a 2.48 Mb inversion specific to chromosome 3 of Sha (Supplementary Fig. 1 and Supplementary Table 12) which explains the suppression of meiotic recombination in this region in hybrids including the Sha haplotype^26,27^. Sequence divergence in rearranged regions was generally higher as compared to collinear regions mostly due an excess of local copy-gain and copy-loss variation in rearranged regions (Fig. 2a, Supplementary Fig. 2 and Supplementary Table 13).

**Fig. 2.**
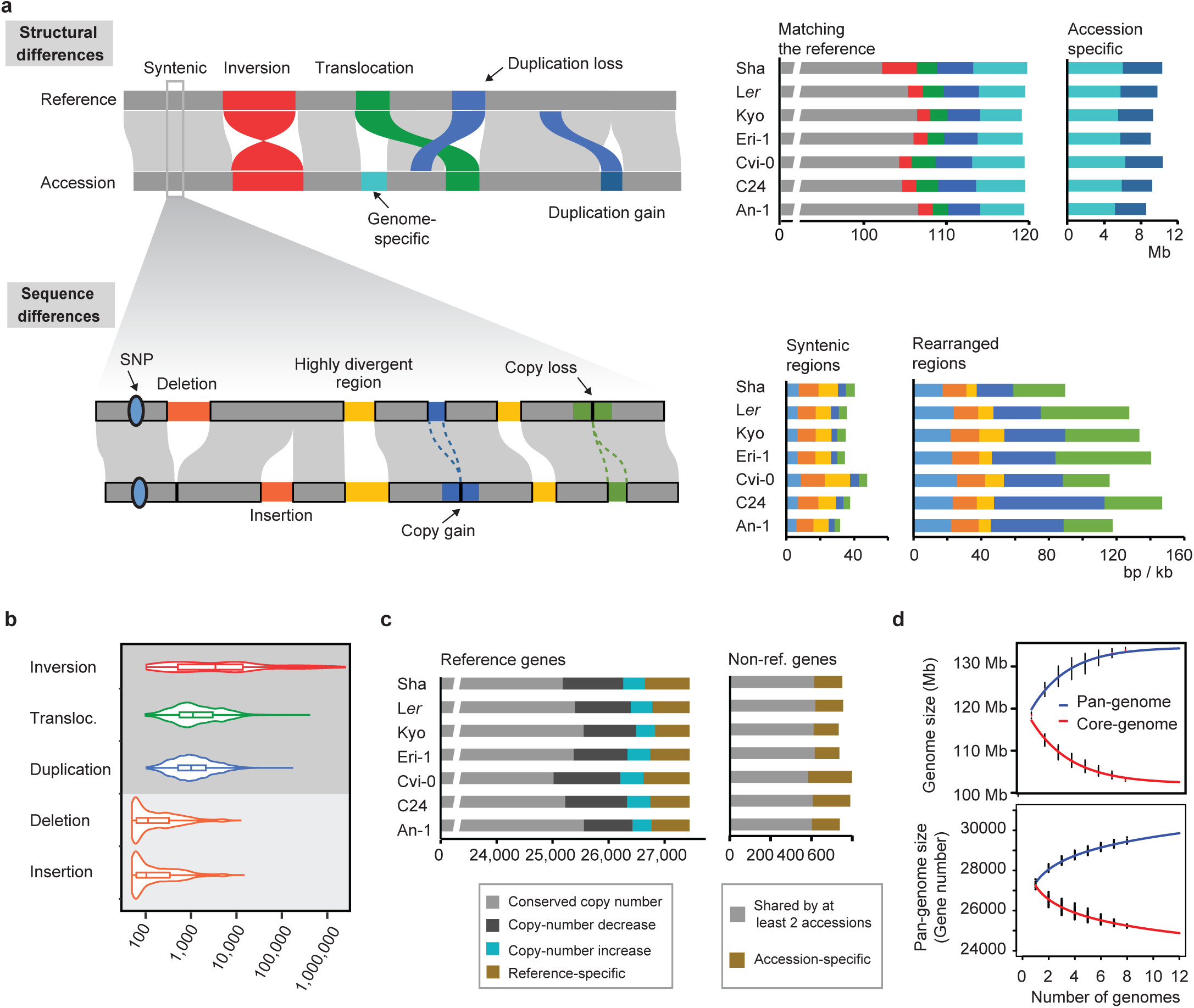
Structural and sequence differences between the genomes. **(a)** Schematic of the structural differences (upper panel) and sequence variation (lower panel) that can be identified between chromosome-level assemblies. Note, local sequence variation can reside in syntenic as well as in rearranged regions. The barplots on the right upper side show the total span of syntenic and rearranged regions between the reference and each of other accessions (colors match the schematic on the left): The left barplots shows the sequence span in respect to the reference sequence, while the right plot shows the sequence space, which is specific to each of the accessions. The barplots on the right lower side show local sequence variation (per kb) in syntenic (left) and rearranged (right) regions between the reference and each of other accessions (again colors match the schematic on the left). **(b)** Size distributions of different types of structural and sequence variation. **(c)** Gene copy number variations between the reference and each of the accessions. The left barplots shows the fraction of (reference) genes which are in gene families with conserved or variable copy numbers. The right barplots shows the number of non-reference genes found in at least two accessions, or found to be specific to an accession genome. **(d)** Pan-genome and core-genome estimations for sequence (upper plot) and gene space (lower plot) were based on all pairwise whole-genome and gene set comparisons across all eight accessions. Each black point corresponds to a pan- or core-genome size estimated with a particular combination of genomes. Pan-genome (blue) and core-genome (red) estimations were fitted using an exponential model.

Genomic rearrangements have the potential to delete, create or duplicate genes resulting in gene copy number variation (CNV). Based on the clustering of orthologous genes across all eight accessions^28^ we found 22,040 gene families with conserved copy number, while 4,957 gene families showed differences in gene copy numbers in at least one accession (Fig. 2c and Supplementary Table 14). Almost 99% of these copy-variable gene families had a maximum copy number of 5 or less, while only less than 10% of them showed more than two different copy numbers across the eight accessions (Supplementary Fig. 3). Among the copy-variable genes we found 1,941 non-reference gene families including 891 gene families present in at least two of the other accessions (Fig. 2c). Around 23% of the non-reference gene families featured orthologs in the closely related genome of *Arabidopsis lyrata* and, according to RNA-seq read mapping, 26%-40% of them showed evidence of expression (Supplementary Table 15). The remaining 1,050 non-reference (accessions-specific) gene families were evenly distributed across the accessions (Fig. 2c), with the exceptions of Cvi-0, where we found nearly twice as many (214) accession-specific genes which is in agreement with the divergent ancestry of this relict accession^12,29^.

**Fig. 3.**
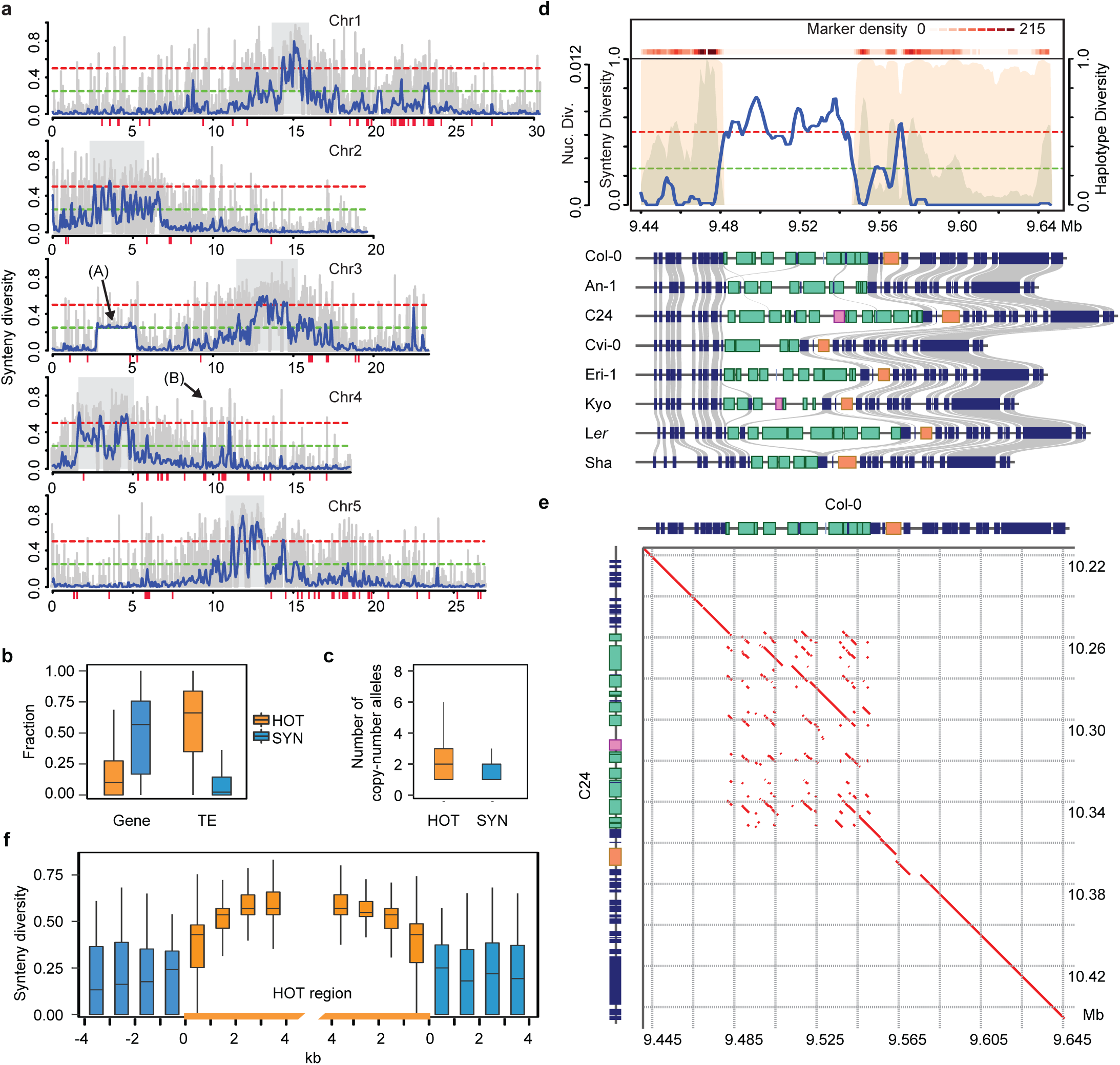
Quantitative analysis of synteny reveals hotspots of rearrangements. **(a)** *Synteny Diversity* along each chromosome: (100kb sliding windows with a step-size of 50kb in blue; 5kb sliding windows with a step-size of 1kb in grey). The red bars under the x-axes indicate the location of R gene clusters. Gray rectangles indicate the location of centromeric regions. The dashed green and red lines indicate thresholds for Synteny Diversity values of 0.25 and 0.50 indicative for the segregation of two (0.25) or three (0.50) non-syntenic haplotypes. The arrow labelled with “(A)” indicates a 2.48 Mb inversion in the Sha genome. The arrow labelled with “(B)” indicates the location of the example show in (b). **(b)** Gene and TE densities in syntenic (SYN) and hotspots of rearrangements (HOT) regions. **(c)** The number of variable copy-number alleles in syntenic (SYN) and rearrangements (HOT) regions**. (d)** An example of a HOT region which includes the *RPP4/RPP5* R gene cluster. The upper panel shows the distribution of *Synteny Diversity* (blue curve) nucleotide diversity (gray background) and haplotype diversity (pink background) in a 5kb sliding window with a step-size of 1kb. Both the nucleotide diversity and the haplotype diversity were calculated based on informative markers (MAF >= 0.05, missing rate < 0.2) from the 1001 Genomes Project ^12^. The marker density is shown as the heatmap on top. The green and red dashed lines indicate the value 0.25 and 0.50 of synteny diversity, respectively. The schematic in the lower part shows the protein-coding genes (colored rectangles) annotated in each of the eight genomes. In blue, genes without function implicated in disease resistance. Rectangles with other colors represent resistance genes, where genes with the same color belong to the same gene family. The gray links between the rectangles indicate the homologous relationships between non-resistance genes. The red lines indicate the positions of HOT regions**. (e)** A dot plot of Col-0 and C24 sequence from the HOT region shown in (D). Red lines indicate regions with homology between the two genomes. **(f)** The distribution of *Synteny Diversity* values in 1kb sliding windows around and in HOT regions.

Based on all possible pairwise genome comparisons, we identified 5.1 - 6.5 Mb accession-specific sequence and used this to estimate a pan-genome size of ∼135 Mb including ∼30,000 genes and a core-genome size of ∼105 Mb with ∼24,000 genes (Fig. 2d)^30,31^ illustrating that one reference genome is not sufficient to capture the entire sequence diversity within *A. thaliana*.

As most of the *A. thaliana* genomes have been analyzed with short read sequencing hardly anything is known about their collinearity. In contract, chromosome-level assemblies allow for a first analysis of the conservation of the contiguity between multiple individuals. To quantify the collinearity we developed a new parameter called *Synteny Diversity π*_*syn*_, which is similar to *Nucleotide Diversity*^32^, however, instead of measuring average sequence differences it measures the degree of collinearity between the genomes of a population (see Methods). *π*_*syn*_ values can range from 0 to 1, where 1 refers to the complete absence of collinearity between any of the genomes and 0 to regions where all genomes are collinear. *π*_*syn*_ can be calculated in any given region, however, the annotation of collinearity still needs to be established within the context of the whole genomes to avoid false assignments of homologous but non-allelic regions.

We calculated *π*_*syn*_ in 5-kb sliding windows across the genome using pair-wise comparisons of all eight accessions (Fig. 3a). As expected, *π*_*syn*_ was generally high in pericentromeric regions and low in chromosome arms. Overall, this revealed around 90 Mb (76% of the genome) where all genomes were collinear to each other, while for the remaining 29 Mb (24%) the collinearity between the genomes was not conserved. This, for example, included a region on chromosome 3 (ranging from Mb ∼2.8 - 5.3), where *π*_*syn*_ was increased to ∼0.25 (i.e. one genome is not collinear to all other seven genomes) due to the 2.48 Mb inversion in the Sha genome (Fig. 3a, arrow labelled with (A)).

Unexpectedly, however, some regions featured *π*_*syn*_ values even larger than 0.5. This implied that not only two, but multiple independent, non-collinear haplotypes segregate in these regions. In turn, this suggest that these regions are more likely to undergo or conserve complex mutations as compared to the rest of the genomes and thereby create *hotspots of rearrangements* (HOT regions) where multiple accessions independently evolved diverse haplotypes. Overall, we found 576 of such HOT regions with a total size of 10.2 Mb including 351 HOT regions in the gene-rich chromosome arms with a total length of 4.1 Mb (or 4% of euchromatic genome) (Supplementary Table 16).

Even though HOT regions in euchromatic regions included more transposable elements and less genes as compared to the collinear regions, they still contained significant numbers of genes, many of which occurred at high and variable copy-number between the accessions (Fig. 3b,c). For example, a HOT region on chromosome 4, which overlapped with the *RPP4/RPP5* R gene cluster^33,34^, displayed 5-15 intact or truncated copies of the *RPP5* gene across the eight genomes (Fig. 3d and Supplementary Table 17). The different gene copies were primarily introduced by an accumulation of forward tandem duplications and large indels (Fig. 3e).

This remarkable pattern of forward tandem duplications and large indels was shared by many of the HOT regions (Fig. 3c and Supplementary Fig. 4). The clear pattern of almost exclusively forward tandem duplications suggested higher mutation (duplication) rates, which are specific to these regions in each of the accessions. In contrast, the borders of the HOT regions were surprisingly well conserved across the accessions (Fig. 3f). This suggested that either different selection regimes introduced clear-cut borders between the HOT regions and their vicinity, or that HOT regions are specific targets of increased tandem duplication rates. Such a local increase of mutation rates could potentially be mediated by non-allelic homologous recombination which could be triggered by the high number of local repeats in these regions^35,36^. Fig. 4 shows two more examples of these complex regions.

**Fig. 4.**
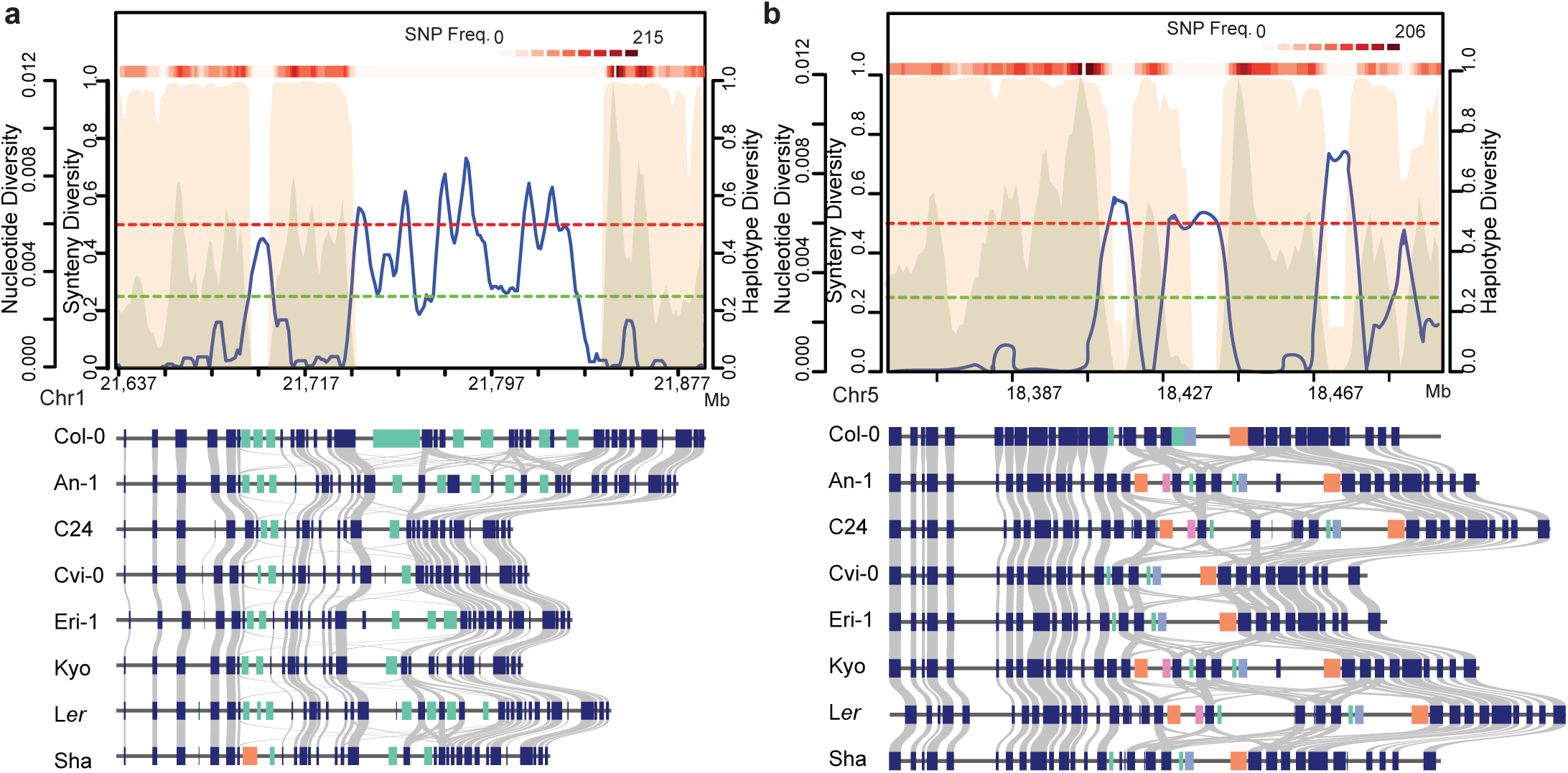
Two examples for hotspots of rearrangements. Visualization of **(a)** the *DM6* locus (*RPP7*) and **(b)** an unnamed R gene cluster on chromosome 5. Descriptions for the plots can be found in the legend of Fig. 3d.

In contrast, meiotic recombination in Arabidopsis was shown to be supressed by structural diversity^37^. To test if HRs are indeed depleted for meiotic recombination, we overlapped rearranged regions with 15,683 crossover (CO) sites previously identified within Col-0/Ler F2 progenies^37,38^. Only 64 of them partially overlapped with non-syntenic regions while all other COs were found in syntenic regions (Fig. 5a), suggesting that HOT regions are almost completely silenced for COs (P-value < 2e-16). In consequence, this would imply that HOT regions are segregating as large non-recombining regions. To test this, we analysed the linkage disequilibrium (LD) within 1,135 genomes of the 1001 Genomes Project^12^ around and across the HOT regions. LD increased in the vicinity of the HOT regions, with increasing LD close to the HOT regions implying reduced recombination in the regions surrounding the HOT regions. Likewise, LD was also high within HOT regions corroborating the recombination suppression in HOT regions. However, when calculated across the border of these regions, LD was significantly lower supporting the idea that HOT regions are not strongly linked to the surrounding haplotypes and that they hardly exchange alleles (Fig. 5b).

**Fig. 5.**
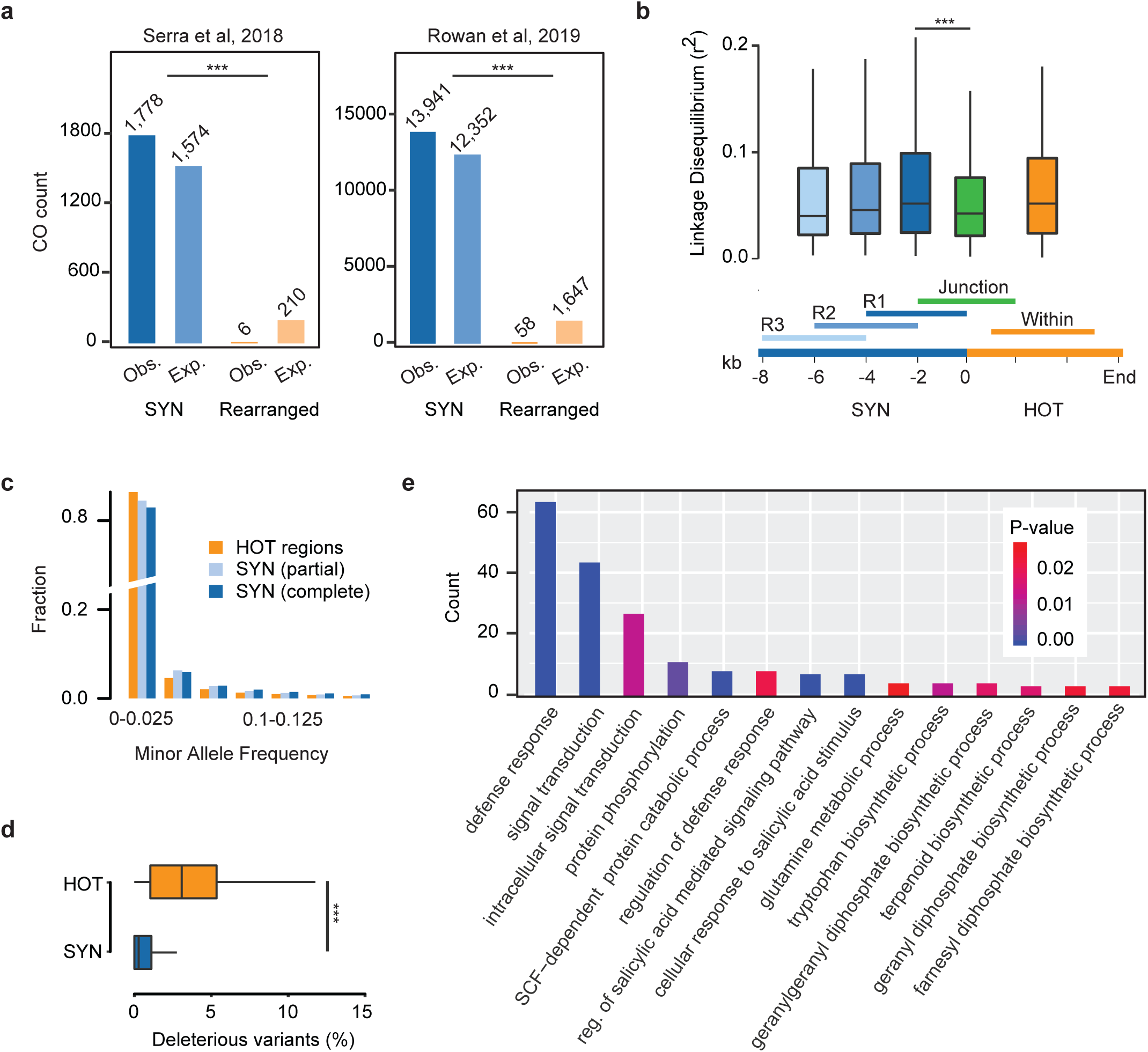
The causes and consequences of hotspots of rearrangements. **(a)** Crossover (CO) breakpoints ^37,38^ identified in Col-0 x L*er* hybrids were checked for their overlaps in syntenic or rearranged regions. Only unique CO intervals smaller than 5kb were used. Chi-square test was applied. **(b)** Linkage disequilibrium (LD) calculated in 4kb windows in and around each of the HOT regions as shown in the lower part. LD was calculated as the correlation coefficient (r^2^) based on informative SNP markers (MAF > 0.05, missing rate < 0.2) selected from the 1001 Genomes Project data ^12^. **(c)** Minor allele frequency of SNP markers in syntenic, partially syntenic, and HOT regions. The SNP markers (MAF >0.005, missing rate < 0.2) from 1001 Genomes Project were used. **(d)** Frequency of deleterious mutations in syntenic regions and HOT regions. Deleterious mutations include SNPs and small indels that introduce premature stop codons, loss of start or stop codons, frameshifts, splicing sites mutations or deletions of exons. (**e**) GO term enrichment analysis of protein-coding genes in HOT regions (P-values cut-off = 0.05).

Reduced meiotic recombination has been linked to the accumulation of new (deleterious) mutations^39^. In agreement with this, HOT regions showed an accumulation of SNPs with low allele frequencies and potentially deleterious variation as compared to other regions in the genome (Fig. 5c,d and Supplementary Fig. 5). Moreover, reduced recombination combined with geographic isolation can provide the basis for the development of alleles, which are incompatible with distantly related haplotypes leading to intra-species incompatibilities^40^. To test this, we searched the location of nine recently reported genetic incompatible loci^41^ (*DM1*-*9*) and found that all except of one overlapped with HOT regions, while *DM3*, the locus which did not overlap with a HOT region, was closely flanked by two HOT regions (Supplementary Fig. 6-12). In addition, we also checked the locus of a recently published single-locus genetic incompatibility^42^ and found that it was also residing in a HOT region (Supplementary Fig. 13).

The high structural diversity of the HOT regions was reminiscent of the patterns that have been described for R gene clusters^43–50^. In fact, the 808 reference genes in HOT regions were significantly enriched for genes involved in defense response, signal transduction and secondary metabolite biosynthesis (Fig. 5e) suggesting a reoccurring role of HOT regions in the adaptation to biotic stress.

As biotic stress is an evolving environmental challenge, the Red Queen hypothesis suggests that the genomes of *A. thaliana* are in the constant need to diversify their offspring^51^. It has been proposed that in response to this, meiotic recombination might increase and thereby diversified offspring is generated^52^. However, exclusively shuffling existing variation might not be sufficient to respond to the evolution of pathogens. Instead, it has been proposed that the accumulation of new gene duplicates could enable a rapid genomic response of plants against pathogens^36,53–55^. The hotspots of rearrangements have the potential to build the basis for such a response, as frequent gene duplications could build the basis for an evolutionary playground to evolve a quick response to the challenges of biotic stress and overcome fitness valleys during the evolution of more complex function. This, in turn, comes at the costs of loss of synteny and the loss of meiotic recombination between distant haplotypes. Though it still needs to be analyzed whether local populations show the same level of diversity or if their haplotypes in HOT regions are more similar and still exchange alleles, we have observed the negative consequences of reduced meiotic recombination in this small world-wide population including the accumulation of deleterious alleles and incompatible epistatic effects between distant genotypes.

Taken together, using chromosome-level genome assemblies of a small, highly diverse population of *A. thaliana*, we have identified regions where genome collinearity (a prerequisite for the allelic exchange in populations) was lost due to genome-specific accumulation of rearrangements. We propose that these regions, which we called *hotspots of rearrangements* or HOT regions, facilitate evolutionary responses to rapidly changing environmental challenges through the accumulation of mutations and not by shuffling alleles. These regions are thus undergoing different evolutionary dynamics as compared to the rest of the genome, where each region segregates with only few haplotypes. Future genome-wide screens for selection patterns might need to take the nature of such regions into account.

## Methods

### Plant material and whole genome sequencing

We received the seeds of all seven accessions from Maarten Koornneef (MPI for Plant Breeding Research), and grew them under normal greenhouse conditions. DNA preparation and next generation sequencing was performed by the Max Planck Genome center. The DNA samples were prepared as previously described for the L*er*^17^, and sequenced with a PacBio Sequel system. For each accession, data from two SMRT cells were generated. Besides, Illumina paired-end libraries were prepared and sequenced on the Illumina HiSeq system.

### Genome assembly

PacBio reads were filtered for short (<50bp) or low quality (QV<80) reads using SMRTLink5 package. *De novo* assembly of each genome was initially performed using three different assembly tools including Falcon^21^, Canu^56^ and MECAT^57^. The resulting assemblies were polished with Arrow from the SMRTLink5 package and then further corrected with mapping of Illumina short reads using BWA^58^ to remove small-scale assembly errors which were identified with SAMTools^59^. For each genome, the final assembly was based on the Falcon assembly as these assemblies always showed highest assembly contiguity. A few contigs were further connected or extended based on whole genome alignments between Falcon and Canu or MECAT assemblies. Contigs were labelled as organellar contigs if they showed alignment identity and coverage both larger than 95% when aligned against the mitochondrial or chloroplast reference sequences. A few of contigs aligned to multiple chromosomes and were split if no Illumina short read alignments supported the conflicting regions. Assembly contigs larger than 20kb were combined to pseudo-chromosomes according to their alignment positions when aligned against the reference sequence using MUMmer4^60^. Contigs with consecutive alignments were concatenated with a stretch of 500 Ns. To note, the assembly of the L*er* accession was already described in a recent study^17^.

### Assembly evaluation

We evaluated the assembly completeness by aligning the reference genes against each of the seven genomes using Blastn^61^. Reference genes which were not aligned or only partially aligned might reveal genes which were missed during the assembly. To examine whether they were really missed, we mapped Illumina short reads from each genome against the reference genome using the BWA^58^ and checked the mapping coverage of these genes. The genes, which were missing in the assembly, should show fully alignment coverage (Supplementary Table 7).

Centromeric and telomeric tandem repeats were annotated by searching for the 178 bp tandem repeat unit^62^ and the 7 bp tandem repeat unit of TTTAGGG^63^. rDNA clusters were annotated with Infernal version 1.1^64^.

The assembly contiguity of Cvi-0 and L*er* were further tested using three previously published genetic maps^65–67^ (Supplementary Table 3). For this we aligned the marker sequences against the chromosome-level assemblies and checked the order of the markers in the assembly versus their order in the genetic map. The ordering of contigs to chromosomes was perfectly supported by all three maps. Overall, only six (out of 1,156) markers showed conflicts between the genetic and physical map. In all six cases we found evidence that the conflict was likely caused by structural differences between the parental genomes.

### Gene annotation

Protein-coding genes were annotated based on *ab initio* gene predications, protein sequence alignments and RNA-seq data. Three *ab initio* gene predication tools were used including Augustus^68^, GlimmerHMM^69^ and SNAP^70^. The reference protein sequences from the Araport 11^24^ annotation were aligned to each genome assembly using exonerate^71^ with the parameter setting “--percent 70 --minintron 10 --maxintron 60000”. For five accessions (An-1, C24, Cvi-0, Ler-0, and Sha) we downloaded a total of 155 RNA-seq data sets from the NCBI SRA database (Supplementary Table 8). RNA-seq reads were mapped to the corresponding genome using HISAT2^72^ and then assembled into transcripts using StringTie^73^ (both with default parameters). All different evidences were integrated into consensus gene models using Evidence Modeler^74^.

The resulting gene models were further evaluated and updated using the Araport 11^24^ annotation. Firstly, for each of the seven genomes, the predicted gene and protein sequences were aligned to the reference sequence, while all reference gene and protein sequences were aligned to each of the other seven genomes using Blast^61^. Then, potentially mis-annotated genes including mis-merged (two or more genes are annotated as a single gene), mis-split (one gene is annotated as two or more genes) and unannotated genes were identified based on the alignments using in-house python scripts. Mis-annotated or unannotated genes were corrected or added by incorporating the open reading frames generated by *ab initio* predications or protein sequence alignment using Scipio^75^.

Noncoding genes were annotated by searching the Rfam database^76^ using Infernal version 1.1^64^. Transposon elements were annotated with RepeatMasker (http://www.repeatmasker.org). Disease resistance genes were annotated using RGAugury77. NB-LRR R gene clusters were defined based on the annotation from a previous study^78^.

### Pan-genome analysis

Pan-genome analyses were performed at both sequence and gene level. To construct a pan-genome of sequences, we generated pair-wise whole genome sequence alignments of all possible pairs of the eight genomes using the nucmer in the software package MUMmer4^60^. A pan-genome was initiated by choosing one of the genomes, followed by iteratively adding the non-aligned sequence of one of the remaining genomes. Here, non-aligned sequences were required to be longer than 100bp without alignment with an identity of more than 90%. The core genome was defined as the sequence space shared by all sampled genomes. Like the pan-genome, the core-genome analysis was initiated with one genome. Then all other genomes were iteratively added, while excluding all those regions which were not aligned against each of the other genomes. The pan- and core-genome of genes was built in a similar way. The pan-genome of genes was constructed by selecting the whole protein coding gene set of one of the accessions followed by iteratively adding the genes of one of the remaining accessions. Likewise, the core-genome of genes was defined as the genes shared in all sampled genomes.

For each pan or core genomes analysis, all possible 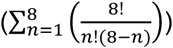 combinations of integrating the eight genomes (or a subset of them) were evaluated. The exponential regression model y = A e^Bx^ + C to was then used to model the pan-genome/core-genomes by fitting medians using the least square method implemented in the nls function of R.

### Analysis of structural rearrangements and gene CNV

All assemblies were aligned to the reference sequence using nucmer from the MUMmer4^60^ toolbox with parameter setting “-max -l 40 -g 90 -b 100 -c 200”. The resulting alignments were further filtered for alignment length (>100) and identity (>90). Structural rearrangements and local variations were identified using SyRI^17^. The functional effects of sequence variation were annotated with snpEff^79^. The gene CNV were identified according to the gene family clustering using the tool OrthoFinder^28^ based on all protein sequences from the eight accessions.

### Synteny diversity, hotspots of rearrangements and diversity estimates

*Synteny Diversity* was defined as the average fraction of non-syntenic sites found within all pairwise genome comparisons within a given population. Here we denote *Synteny Diversity* as

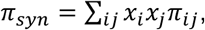

where *x*_*i*_ and *x*_*j*_ refer to the frequencies of sequence *i* and _*j*_ and *π*_*ij*_ to the average probability of a position in *i* to be non-syntenic. Note, *π*_*syn*_ can be calculated in a given region or for the entire genome. However even when calculated for small regions the annotation of synteny still needs to established within the context of the whole genomes to avoid false assignments of homologous but non-allelic sequence. Here we used the annotation of *SyRI* to define syntenic regions. *π*_*syn*_ values can range from 0 to 1, with higher values referring to a higher average degree of non-syntenic regions between the genomes.

For the analyses, we calculated *π*_*syn*_ in 5-kb sliding windows with 1kb step-size across the entire genome. HOT regions were defined as regions with *π*_*syn*_ larger than 0.5. Neighboring regions were merged into one HOT region if their distance was shorter than 2kb.

The nucleotide and haplotype diversity were calculated with the R package PopGenome^80^ using SNP markers (with MAF > 0.05) from 1001 Genomes Project^12^. LD were calculated as correlation coefficients *r*^2^ using SNP markers with MAF > 0.05. GO enrichment analysis was performed using the webtool DAVID^81,82^

## Data availability

Raw sequencing data, assemblies and annotations can be accessed in the European Nucleotide Archive under the project accession number PRJEB31147. Assemblies, annotation, variation and orthologs can be found on the *1001 Arabidopsis thaliana Genomes* webpage (www.1001genomes.org).

## Code availability

Custom code used in this study can be found online at https://github.com/schneebergerlab/AMPRIL-genomes.

## Supplementary Information

Tables S1 to S17

Figs. S1 to S15

## Acknowledgements

The authors would like to thank Beth R. Rowan (UC Davis) for providing the CO breakpoint list prior to publication, Bruno Hüttel (Max Planck Genome center) for support in genome sequencing, Sigi Effgen and Maarten Koornneef (Max Planck Institute for Plant Breeding Research) for providing seeds, Onur Dogan (Max Planck Institute for Plant Breeding Research) for help in the greenhouse, Angela M. Hancock (Max Planck Institute for Plant Breeding Research) for helpful discussions, and Raphael Mercier and Padraic J. Flood (Max Planck Institute for Plant Breeding Research) for helpful comments on the manuscript and the interpretation of HOT regions.

## Authors contributions

W.-B.J. and K.S. designed the study. W.-B.J. performed all analysis. K.S. supervised the study. W.-B.J. and K.S. wrote the manuscript. All authors read and approved the final manuscript.

