## Supplementary Tables and Figures for "Chromosome-level assemblies of multiple Arabidopsis genomes reveal hotspots of rearrangements with altered evolutionary dynamics"

Jiao and Schneeberger, 2019

.

### Supplementary Tables

#### Supplementary Table 1. Whole genome DNA sequencing data.

| **Accession** |  | **PacBio** | | |  | **Illumina paired-end** | | |
| --- | --- | --- | --- | --- | --- | --- | --- | --- |
|  |  | **Reads** | **Mean Length** | **Depth** |  | **Pairs** | **Length** | **Depth** |
| An-1 |  | 1,419,095 | 6,745 | 70 |  | 52,408,462 | 101 | 78.4 |
| C24 |  | 1,026,041 | 6,011 | 45 |  | 37,543,070 | 101 | 56.2 |
| Cvi-0 |  | 1,358,030 | 7,132 | 71 |  | 44,119,747 | 101 | 66.0 |
| Eri-1 |  | 1,364,672 | 6,402 | 64 |  | 37,489,627 | 101 | 56.1 |
| L*er* |  | 1,312,624 | 7,140 | 69 |  | 48,030,713 | 101 | 71.9 |
| Kyo |  | 1,254,045 | 6,599 | 61 |  | 42,068,024 | 101 | 62.9 |
| Sha |  | 853,955 | 8,281 | 52 |  | 50,271,422 | 101 | 75.2 |

#### **Supplementary Table** **2. Contig assembly statistics.**

|  | **An-1** | **C24** | **Cvi-0** | **Eri-1** | **Kyo** | **L*er*** | **Sha** |
| --- | --- | --- | --- | --- | --- | --- | --- |
| **Contigs** | 151 | 167 | 140 | 200 | 230 | 149 | 143 |
| **Total bases** | 120,109,838 | 119,191,266 | 119,730,512 | 120,766,191 | 122,179,079 | 120,316,059 | 120,265,401 |
| **N25** | 11,780,999 | 9,748,094 | 9,283,899 | 8,805,940 | 11,196,314 | 13,234,969 | 12,200,080 |
| **N50** | 8,166,377 | 4,802,170 | 7,363,394 | 4,757,538 | 9,149,088 | 11,168,341 | 6,996,577 |
| **N75** | 4,989,922 | 2,108,767 | 5,593,362 | 1,771,097 | 3,917,708 | 6,074,025 | 2,595,543 |
| **N90** | 1,333,704 | 593,800 | 1,975,628 | 657,186 | 949,167 | 804,247 | 1,088,930 |
| **L25** | 3 | 3 | 3 | 3 | 3 | 3 | 3 |
| **L50** | 6 | 8 | 7 | 8 | 6 | 5 | 6 |
| **L75** | 10 | 18 | 11 | 18 | 10 | 9 | 12 |
| **L90** | 17 | 34 | 17 | 34 | 19 | 18 | 23 |
| **CN50** | 8,166,377 | 4,802,170 | 6,994,145 | 4,757,538 | 9,149,088 | 13,234,969 | 12,200,080 |
| **CL50** | 2 | 2 | 2 | 2 | 2 | 1 | 1 |
| **NG25** | 11,780,999 | 8,058,848 | 9,238,557 | 7,925,394 | 11,196,314 | 13,234,969 | 12,200,080 |
| **NG50** | 7,186,630 | 3,510,498 | 6,994,145 | 4,279,918 | 9,149,088 | 8,759,918 | 5,319,415 |
| **NG75** | 2,765,233 | 1,004,113 | 2,403,042 | 1,106,604 | 2,177,877 | 2,307,265 | 1,494,253 |
| **NG90** | 0 | 0 | 0 | 0 | 12,670 | 0 | 0 |
| **LG25** | 3 | 4 | 4 | 4 | 3 | 3 | 3 |
| **LG50** | 7 | 10 | 8 | 10 | 6 | 6 | 7 |
| **LG75** | 13 | 26 | 15 | 25 | 13 | 12 | 18 |
| **LG90** | 0 | 0 | 0 | 0 | 132 | 0 | 0 |
| **Min** | 508 | 86 | 488 | 561 | 474 | 184 | 1,016 |
| **Max** | 14,839,891 | 10,639,543 | 12,641,456 | 12,399,038 | 14,879,663 | 14,816,562 | 14,310,783 |

#### Supplementary Table 3. Assembly validation using genetic maps.

| **Cross** | **Assembly tested** | **Data** | **Markers** | **Aligned** | **Conflicts** |
| --- | --- | --- | --- | --- | --- |
| Cvi-0 x Col-0 | Cvi-0 | Simon et al., 2008^1^ | 94 | 93 | 0 |
| L*er* x Col-0 | L*er* | Singer et al., 2006^2^ | 676 | 415 | 3 |
| L*er* x Col-0 | L*er* | Giraut et al., 2011^3^ | 386 | 357 | 3 |

**Note**: Three genetic maps were used to validate the assemblies of Cvi-0 and L*er*. Evaluation of the conflicting markers revealed that all them of were likely caused by structural variations. Each of the conflicting markers was found on contigs which were supported by non-conflicting markers as well.

#### Supplementary Table 4. Location of centromeric and telomeric tandem repeat arrays.

See attached Supplementary.Tables.v1.xlsx files.

#### Supplementary Table 5. Location of rDNA clusters.

| **Accession** | **Chr** | **Start** | **End** | **Size** | **Unit Count** |
| --- | --- | --- | --- | --- | --- |
| An-1 | chr1 | 16,247,697 | 16,249,746 | 2,049 | 5 |
| An-1 | chr2 | 15,160 | 30,883 | 15,723 | 4 |
| An-1 | chr3 | 13,660,309 | 13,693,016 | 32,707 | 44 |
| An-1 | chr4 | 303 | 12,836 | 12,533 | 3 |
| An-1 | chr5 | 11,704,997 | 11,712,070 | 7,073 | 12 |
| An-1 | chr5 | 12,502,040 | 12,517,915 | 15,875 | 41 |
| An-1 | chr5 | 12,528,355 | 12,548,970 | 20,615 | 56 |
| C24 | chr1 | 15,480,421 | 15,481,987 | 1,566 | 4 |
| C24 | chr2 | 5,277 | 9,979 | 4,702 | 3 |
| C24 | chr3 | 14,477,578 | 14,508,366 | 30,788 | 40 |
| C24 | chr4 | 3,024,558 | 3,029,149 | 4,591 | 10 |
| C24 | chr5 | 11,167,024 | 11,173,642 | 6,618 | 11 |
| C24 | chr5 | 12,047,791 | 12,084,720 | 36,929 | 86 |
| Cvi-0 | chr1 | 15,291,551 | 15,293,036 | 1,485 | 1 |
| Cvi-0 | chr1 | 16,201,388 | 16,203,919 | 2,531 | 6 |
| Cvi-0 | chr3 | 13,553,363 | 13,557,056 | 3,693 | 8 |
| Cvi-0 | chr3 | 13,632,106 | 13,672,892 | 40,786 | 50 |
| Cvi-0 | chr3 | 13,693,322 | 13,720,728 | 27,406 | 49 |
| Cvi-0 | chr3 | 14,005,932 | 14,048,548 | 42,616 | 37 |
| Cvi-0 | chr4 | 3,078,679 | 3,107,071 | 28,392 | 57 |
| Cvi-0 | chr5 | 11,309,985 | 11,316,573 | 6,588 | 11 |
| Eri-1 | chr1 | 15,214,182 | 15,216,702 | 2,520 | 6 |
| Eri-1 | chr3 | 14,398,975 | 14,411,194 | 12,219 | 25 |
| Eri-1 | chr3 | 14,944,897 | 14,945,898 | 1,001 | 1 |
| Eri-1 | chr5 | 12,503,702 | 12,518,335 | 14,633 | 37 |
| Eri-1 | chr5 | 12,528,934 | 12,547,045 | 18,111 | 48 |
| Kyo | chr1 | 15,615,850 | 15,617,896 | 2,046 | 5 |
| Kyo | chr3 | 14,537,056 | 14,550,096 | 13,040 | 9 |
| Kyo | chr3 | 15,246,489 | 15,247,490 | 1,001 | 1 |
| Kyo | chr5 | 11,290,583 | 11,292,705 | 2,122 | 5 |
| Kyo | chr5 | 11,328,618 | 11,335,208 | 6,590 | 11 |
| Kyo | chr5 | 12,294,721 | 12,301,528 | 6,807 | 15 |
| Kyo | chr5 | 12,324,038 | 12,325,865 | 1,827 | 4 |
| L*er* | chr1 | 16,112,353 | 16,114,394 | 2,041 | 5 |
| L*er* | chr2 | 24,660 | 34,941 | 10,281 | 2 |
| L*er* | chr3 | 6,075,317 | 6,076,940 | 1,623 | 4 |
| L*er* | chr3 | 14,214,467 | 14,253,261 | 38,794 | 7 |
| L*er* | chr3 | 14,267,148 | 14,295,446 | 28,298 | 39 |
| L*er* | chr4 | 4,789 | 31,377 | 26,588 | 6 |
| L*er* | chr4 | 3,461,906 | 3,478,278 | 16,372 | 31 |
| L*er* | chr5 | 11,240,780 | 11,255,952 | 15,172 | 31 |
| L*er* | chr5 | 11,317,853 | 11,324,980 | 7,127 | 12 |
| L*er* | chr5 | 12,403,488 | 12,410,158 | 6,670 | 18 |
| L*er* | chr5 | 12,432,957 | 12,436,284 | 3,327 | 7 |
| Sha | chr1 | 16,317,928 | 16,319,976 | 2,048 | 5 |
| Sha | chr2 | 4,747 | 17,222 | 12,475 | 3 |
| Sha | chr3 | 13,328,855 | 13,413,311 | 84,456 | 37 |
| Sha | chr4 | 1,132 | 27,909 | 26,777 | 5 |
| Sha | chr4 | 39,104 | 50,775 | 11,671 | 5 |
| Sha | chr4 | 2,996,768 | 3,020,817 | 24,049 | 48 |
| Sha | chr5 | 11,301,764 | 11,362,233 | 60,469 | 121 |
| Sha | chr5 | 12,959,621 | 12,973,107 | 13,486 | 34 |

#### Supplementary Table 6. Sequence annotation near assembly gaps.

| **Assembly** | **Gaps** | **Pericentromere** | **rDNA** | **TE** | **All** |
| --- | --- | --- | --- | --- | --- |
| An-1 | 40 | 26 | 2 | 38 | 95.00% |
| C24 | 68 | 34 | 3 | 59 | 86.76% |
| Cvi-0 | 38 | 25 | 6 | 35 | 92.11% |
| Eri-1 | 58 | 22 | 2 | 49 | 84.48% |
| Kyo | 46 | 31 | 3 | 41 | 89.13% |
| L*er* | 44 | 27 | 4 | 41 | 93.18% |
| Sha | 49 | 30 | 5 | 47 | 97.96% |

**Note**: Assembly gaps can reside in the pericentromere, in or close to ribosomal DNA cluster (distance to the closest rDNA cluster < 5kb) or in or close to transposable elements (distance to closest TE < 500bp). As some of them may simultaneously locate in the pericentromere and rDNA clusters or TEs, the last column indicates the percent of gaps that matched at least one of these features.

#### Supplementary Table 7. Assembly evaluation.

| **Accession** | **Genes without any blastn hit**  **(fully deleted)** | | **Genes without high quality blastn hit (partially deleted)** | | **Percent of genes with good blastn hit** | | **No. of genes without full (<60%) short read coverage** |
| --- | --- | --- | --- | --- | --- | --- | --- |
| **An-1** | 112 | 75 | | 99.3% | | 92.5% | |
| **C24** | 120 | 88 | | 99.2% | | 85.6% | |
| **Cvi-0** | 136 | 95 | | 99.2% | | 87.4% | |
| **Eri-1** | 102 | 76 | | 99.4% | | 88.2% | |
| **Kyo** | 85 | 79 | | 99.4% | | 87.8% | |
| **L*er*** | 107 | 74 | | 99.3% | | 89.0% | |
| **Sha** | 159 | 89 | | 99.1% | | 75.8% | |

**Note:** To check the assembly completeness, we aligned all 27,445 reference genes against each assembly using blastn**.** Genes without any blastn hit are fully absent in the assembly (second column). Genes without high-quality blastn hit are only partially present in the assembly (third column). (“High-quality blastn hits” were defined as blastn hits with identity larger than 85 and coverage larger than 95%.)

In order to check if the genes that are absent in the assemblies were just not assembled or were truly missing in the genomes, we further aligned the Illumina whole-genome reads from each accession against Col-0 reference sequence. The last column indicates how many of the genes without blastn hit or good blastn hit have short read alignment coverage in less than 60% of their sequence (and are thus mostly like truly absent in the sequenced genomes).

#### Supplementary Table 8. RNA-seq data used for gene annotations.

| **Accession** | **NCBI/EBI Project ID** | **NCBI/EBI Sample ID** | **Total bases** | **Reference** |
| --- | --- | --- | --- | --- |
| An-1 | ERP009505 | ERS657074 | 6,636,065,822 | Clauw et al, 2015^4^ |
| An-1 | ERP009505 | ERS657079 | 4,807,026,522 | Clauw et al, 2015^4^ |
| An-1 | ERP009505 | ERS657082 | 5,619,041,878 | Clauw et al, 2015^4^ |
| An-1 | ERP009505 | ERS657083 | 3,992,566,562 | Clauw et al, 2015^4^ |
| An-1 | ERP009505 | ERS657090 | 3,959,926,392 | Clauw et al, 2015^4^ |
| An-1 | ERP009505 | ERS657094 | 3,519,597,096 | Clauw et al, 2015^4^ |
| An-1 | ERP016867 | ERS1294514 | 844,772,160 | Clauw et al, 2016^5^ |
| An-1 | ERP016867 | ERS1294515 | 1,493,366,394 | Clauw et al, 2016^5^ |
| C24 | ERP015703 | ERS1164812 | 1,519,136,000 | van Veen et al., 2016^6^ |
| C24 | ERP015703 | ERS1164813 | 1,864,829,750 | van Veen et al., 2016^6^ |
| C24 | ERP015703 | ERS1164814 | 1,882,564,700 | van Veen et al., 2016^6^ |
| C24 | ERP015703 | ERS1164815 | 1,664,880,550 | van Veen et al., 2016^6^ |
| C24 | ERP015703 | ERS1164816 | 1,819,241,200 | van Veen et al., 2016^6^ |
| C24 | ERP015703 | ERS1164817 | 1,729,345,700 | van Veen et al., 2016^6^ |
| C24 | ERP016867 | ERS1294528 | 1,825,005,828 | Clauw et al, 2016^5^ |
| C24 | ERP016867 | ERS1294529 | 1,611,268,806 | Clauw et al, 2016^5^ |
| C24 | SRP026082 | SRS448007 | 3,377,368,997 | Cui et al., 2014^7^ |
| C24 | SRP026082 | SRS448016 | 3,348,840,739 | Cui et al., 2014^7^ |
| C24 | SRP031847 | SRS493101 | 2,762,373,684 | Miller et al., 2015^8^ |
| C24 | SRP031847 | SRS493105 | 2,532,505,080 | Miller et al., 2015^8^ |
| C24 | SRP031847 | SRS493109 | 2,585,474,808 | Miller et al., 2015^8^ |
| C24 | SRP035234 | SRS527275 | 3,382,105,190 | Ding et al., 2014^9^ |
| C24 | SRP035234 | SRS527276 | 2,622,812,844 | Ding et al., 2014^9^ |
| C24 | SRP035234 | SRS527277 | 2,386,391,640 | Ding et al., 2014^9^ |
| C24 | SRP035234 | SRS527278 | 2,967,903,180 | Ding et al., 2014^9^ |
| C24 | SRP042060 | SRS609809 | 3,279,324,400 | Zhao et al., 2014^10^ |
| C24 | SRP042060 | SRS609809 | 3,578,046,400 | Zhao et al., 2014^10^ |
| C24 | SRP042060 | SRS609809 | 3,273,046,400 | Zhao et al., 2014^10^ |
| C24 | SRP045478 | SRS682670 | 1,843,542,294 | Perez-Santángelo et al., 2014^11^ |
| C24 | SRP045478 | SRS682671 | 1,843,542,294 | Perez-Santángelo et al., 2014^11^ |
| C24 | SRP045478 | SRS682672 | 4,032,844,150 | Perez-Santángelo et al., 2014^11^ |
| C24 | SRP045478 | SRS682673 | 4,032,844,150 | Perez-Santángelo et al., 2014^11^ |
| C24 | SRP045478 | SRS682674 | 2,839,123,332 | Perez-Santángelo et al., 2014^11^ |
| C24 | SRP045478 | SRS682675 | 2,839,123,332 | Perez-Santángelo et al., 2014^11^ |
| C24 | SRP047297 | SRS703918 | 1,102,642,440 | Xu et al.,2015^12^ |
| C24 | SRP047297 | SRS703920 | 1,229,302,317 | Xu et al.,2015^12^ |
| C24 | SRP047297 | SRS703919 | 1,160,182,272 | Xu et al.,2015^12^ |
| C24 | SRP047297 | SRS703921 | 1,445,841,024 | Xu et al.,2015^12^ |
| C24 | SRP047297 | SRS703923 | 922,092,699 | Xu et al.,2015^12^ |
| C24 | SRP047297 | SRS703922 | 1,268,096,232 | Xu et al.,2015^12^ |
| C24 | SRP047445 | SRS708049 | 3,382,105,190 | Cui et al., 2016^13^ |
| C24 | SRP047445 | SRS708050 | 2,967,903,180 | Cui et al., 2016^13^ |
| C24 | SRP050410 | SRS775831 | 2,504,507,649 | Du et al., 2016^14^ |
| C24 | SRP051513 | SRS803270 | 6,723,106,600 | Groszmann et al.,2015^15^ |
| C24 | SRP051513 | SRS803271 | 6,815,633,200 | Groszmann et al.,2015^15^ |
| C24 | SRP051513 | SRS803272 | 7,251,068,800 | Groszmann et al.,2015^15^ |
| C24 | SRP051513 | SRS803273 | 7,092,479,200 | Groszmann et al.,2015^15^ |
| C24 | SRP051513 | SRS803274 | 4,925,493,058 | Groszmann et al.,2015^15^ |
| C24 | SRP051513 | SRS803275 | 4,412,760,498 | Groszmann et al.,2015^15^ |
| C24 | SRP051763 | SRS973510 | 7,251,068,800 | Wang et al., 2015^16^ |
| C24 | SRP051763 | SRS973509 | 7,092,479,200 | Wang et al., 2015^16^ |
| C24 | SRP051763 | SRS973489 | 6,532,389,400 | Wang et al., 2015^16^ |
| C24 | SRP051763 | SRS973488 | 5,478,994,000 | Wang et al., 2015^16^ |
| C24 | SRP051763 | SRS810590 | 9,646,799,870 | Wang et al., 2015^16^ |
| C24 | SRP051763 | SRS810591 | 9,812,577,028 | Wang et al., 2015^16^ |
| C24 | SRP051763 | SRS810592 | 8,875,798,190 | Wang et al., 2015^16^ |
| C24 | SRP051763 | SRS810593 | 5,699,734,212 | Wang et al., 2015^16^ |
| C24 | SRP051763 | SRS810618 | 4,925,493,058 | Wang et al., 2015^16^ |
| C24 | SRP051763 | SRS810619 | 4,412,760,498 | Wang et al., 2015^16^ |
| C24 | SRP051764 | SRS810628 | 6,877,736,800 | Wang et al., 2015^16^ |
| C24 | SRP051764 | SRS810629 | 6,367,929,200 | Wang et al., 2015^16^ |
| C24 | SRP052029 | SRS816315 | 733,317,218 | Piofczyk et al., 2015^17^ |
| C24 | SRP052029 | SRS816314 | 1,134,222,706 | Piofczyk et al., 2015^17^ |
| C24 | SRP052029 | SRS816316 | 2,424,693,528 | Piofczyk et al., 2015^17^ |
| C24 | SRP052029 | SRS816318 | 808,021,650 | Piofczyk et al., 2015^17^ |
| C24 | SRP065557 | SRS1142558 | 4,618,230,300 | Zhang et al.,2016^18^ |
| C24 | SRP065557 | SRS1142557 | 5,097,646,952 | Zhang et al.,2016^18^ |
| C24 | SRP065557 | SRS1142556 | 4,555,494,910 | Zhang et al.,2016^18^ |
| C24 | SRP071815 | SRS1345560 | 4,327,166,800 | Zhang et al.,2016^18^ |
| C24 | SRP071815 | SRS1345559 | 4,399,449,600 | Zhang et al.,2016^18^ |
| C24 | SRP071815 | SRS1345558 | 4,733,992,600 | Zhang et al.,2016^18^ |
| C24 | SRP074107 | SRS1415702 | 6,857,983,000 | Kawakatsu et al., 2016^19^ |
| C24 | SRP098906 | SRS1959174 | 4,709,389,800 | Wang et al.,2017^20^ |
| C24 | SRP098906 | SRS1959175 | 4,510,190,600 | Wang et al.,2017^20^ |
| C24 | SRP098906 | SRS1959176 | 4,525,940,800 | Wang et al.,2017^20^ |
| Cvi-0 | ERP009505 | ERS657062 | 3,785,407,280 | Clauw et al, 2015^4^ |
| Cvi-0 | ERP009505 | ERS657068 | 3,716,729,300 | Clauw et al, 2015^4^ |
| Cvi-0 | ERP009505 | ERS657076 | 4,598,817,446 | Clauw et al, 2015^4^ |
| Cvi-0 | ERP009505 | ERS657084 | 2,903,584,966 | Clauw et al, 2015^4^ |
| Cvi-0 | ERP009505 | ERS657086 | 3,788,874,610 | Clauw et al, 2015^4^ |
| Cvi-0 | ERP009505 | ERS657089 | 2,915,870,202 | Clauw et al, 2015^4^ |
| Cvi-0 | ERP015703 | ERS1164824 | 1,831,923,900 | van Veen et al., 2016^6^ |
| Cvi-0 | ERP015703 | ERS1164825 | 1,648,307,850 | van Veen et al., 2016^6^ |
| Cvi-0 | ERP015703 | ERS1164826 | 1,875,332,550 | van Veen et al., 2016^6^ |
| Cvi-0 | ERP015703 | ERS1164827 | 1,858,185,700 | van Veen et al., 2016^6^ |
| Cvi-0 | ERP015703 | ERS1164828 | 1,706,118,650 | van Veen et al., 2016^6^ |
| Cvi-0 | ERP015703 | ERS1164829 | 1,748,830,400 | van Veen et al., 2016^6^ |
| Cvi-0 | SRP074107 | SRS1415705 | 4,915,150,400 | Kawakatsu et al., 2016^19^ |
| Cvi-0 | SRP078926 | SRS1570177 | 2,308,716,250 | Leydon et al., 2017^21^ |
| Cvi-0 | SRP078926 | SRS1570178 | 2,347,827,300 | Leydon et al., 2017^21^ |
| Cvi-0 | SRP078926 | SRS1570214 | 2,623,598,650 | Leydon et al., 2017^21^ |
| Cvi-0 | SRP097638 | SRS1935483 | 455,343,440 | Picard et al., 2017^22^ |
| Cvi-0 | SRP097638 | SRS1935484 | 562,727,560 | Picard et al., 2017^22^ |
| Cvi-0 | SRP097638 | SRS1935485 | 474,340,840 | Picard et al., 2017^22^ |
| Sha | ERP009505 | ERS657064 | 4,143,085,650 | Clauw et al, 2015^4^ |
| Sha | ERP009505 | ERS657067 | 2,596,846,148 | Clauw et al, 2015^4^ |
| Sha | ERP009505 | ERS657081 | 7,357,376,108 | Clauw et al, 2015^4^ |
| Sha | ERP009505 | ERS657085 | 5,783,518,964 | Clauw et al, 2015^4^ |
| Sha | ERP009505 | ERS657088 | 4,809,863,410 | Clauw et al, 2015^4^ |
| Sha | ERP009505 | ERS657095 | 2,964,615,832 | Clauw et al, 2015^4^ |
| Sha | ERP016867 | ERS1294648 | 1,249,672,686 | Clauw et al, 2016^5^ |
| Sha | ERP016867 | ERS1294649 | 1,551,513,534 | Clauw et al, 2016^5^ |
| Sha | SRP074107 | SRS1415556 | 5,320,965,400 | Kawakatsu et al., 2016^19^ |
| Ler | ERP016867 | ERS1294588 | 1,188,327,234 | Clauw et al, 2016^5^ |
| Ler | ERP016867 | ERS1294589 | 1,384,778,520 | Clauw et al, 2016^5^ |
| Ler | SRP051513 | SRS803276 | 7,015,886,800 | Groszmann et al.,2015^15^ |
| Ler | SRP051513 | SRS803277 | 6,661,630,600 | Groszmann et al.,2015^15^ |
| Ler | SRP051513 | SRS803278 | 7,171,574,600 | Groszmann et al.,2015^15^ |
| Ler | SRP051513 | SRS803279 | 6,834,478,800 | Groszmann et al.,2015^15^ |
| Ler | SRP051513 | SRS803280 | 8,238,715,844 | Groszmann et al.,2015^15^ |
| Ler | SRP051513 | SRS803281 | 3,847,275,032 | Groszmann et al.,2015^15^ |
| Ler | SRP051763 | SRS973508 | 7,171,574,600 | Wang et al., 2015^16^ |
| Ler | SRP051763 | SRS973507 | 6,834,478,800 | Wang et al., 2015^16^ |
| Ler | SRP051763 | SRS973487 | 7,302,101,200 | Wang et al., 2015^16^ |
| Ler | SRP051763 | SRS973484 | 5,929,661,000 | Wang et al., 2015^16^ |
| Ler | SRP051763 | SRS810594 | 8,262,935,442 | Wang et al., 2015^16^ |
| Ler | SRP051763 | SRS810595 | 12,625,063,428 | Wang et al., 2015^16^ |
| Ler | SRP051763 | SRS810596 | 7,849,864,228 | Wang et al., 2015^16^ |
| Ler | SRP051763 | SRS810597 | 5,185,351,312 | Wang et al., 2015^16^ |
| Ler | SRP051763 | SRS810620 | 8,238,715,844 | Wang et al., 2015^16^ |
| Ler | SRP051763 | SRS810621 | 3,847,275,032 | Wang et al., 2015^16^ |
| Ler | SRP051764 | SRS810630 | 5,750,736,800 | Wang et al., 2015^16^ |
| Ler | SRP051764 | SRS810631 | 6,262,793,200 | Wang et al., 2015^16^ |
| Ler | SRP058781 | SRS947619 | 1,467,101,700 | Li et al., 2015^23^ |
| Ler | SRP058781 | SRS947619 | 1,179,804,600 | Li et al., 2015^23^ |
| Ler | SRP058781 | SRS947619 | 1,757,773,100 | Li et al., 2015^23^ |
| Ler | SRP058781 | SRS947619 | 1,701,903,600 | Li et al., 2015^23^ |
| Ler | SRP058920 | SRS950133 | 1,593,940,590 | Li et al., 2016^24^ |
| Ler | SRP058920 | SRS950131 | 1,407,251,180 | Li et al., 2016^24^ |
| Ler | SRP065316 | SRS1135750 | 11,354,868,945 | Wuest et al., 2016^25^ |
| Ler | SRP065316 | SRS1135749 | 5,019,108,443 | Wuest et al., 2016^25^ |
| Ler | SRP065316 | SRS1135748 | 3,778,271,125 | Wuest et al., 2016^25^ |
| Ler | SRP074486 | SRS1424474 | 1,769,193,467 | Huang et al., 2016^26^ |
| Ler | SRP074486 | SRS1424476 | 1,534,016,280 | Huang et al., 2016^26^ |
| Ler | SRP074486 | SRS1424477 | 2,052,698,851 | Huang et al., 2016^26^ |
| Ler | SRP083946 | SRS1662551 | 5,937,938,400 | Ezer et al., 2017^27^ |
| Ler | SRP083946 | SRS1662561 | 2,904,970,200 | Ezer et al., 2017^27^ |
| Ler | SRP083946 | SRS1662572 | 6,947,251,800 | Ezer et al., 2017^27^ |
| Ler | SRP083946 | SRS1662592 | 4,842,299,200 | Ezer et al., 2017^27^ |
| Ler | SRP083946 | SRS1662618 | 5,926,146,400 | Ezer et al., 2017^27^ |
| Ler | SRP083946 | SRS1662657 | 6,015,868,400 | Ezer et al., 2017^27^ |
| Ler | SRP083946 | SRS1662679 | 4,986,165,200 | Ezer et al., 2017^27^ |
| Ler | SRP083946 | SRS1662681 | 7,764,758,600 | Ezer et al., 2017^27^ |
| Ler | SRP083946 | SRS1662683 | 2,228,587,800 | Ezer et al., 2017^27^ |
| Ler | SRP083946 | SRS1662686 | 8,868,225,000 | Ezer et al., 2017^27^ |
| Ler | SRP083946 | SRS1662682 | 3,894,951,600 | Ezer et al., 2017^27^ |
| Ler | SRP083946 | SRS1662684 | 2,685,190,800 | Ezer et al., 2017^27^ |
| Ler | SRP083946 | SRS1662688 | 3,794,103,800 | Ezer et al., 2017^27^ |
| Ler | SRP083946 | SRS1662687 | 3,885,968,200 | Ezer et al., 2017^27^ |
| Ler | SRP098906 | SRS1959177 | 4,381,460,200 | Wang et al.,2017^20^ |
| Ler | SRP098906 | SRS1959178 | 4,649,548,400 | Wang et al.,2017^20^ |
| Ler | SRP098906 | SRS1959179 | 4,553,470,400 | Wang et al.,2017^20^ |
| Ler | SRP033660 | SRS513491 | 2,389,096,240 | Li et al., 2017^28^ |
| Ler | SRP033660 | SRS513492 | 2,740,051,484 | Li et al., 2017^28^ |
| Ler | SRP033660 | SRS513493 | 2,090,697,666 | Li et al., 2017^28^ |

#### Supplementary Table 9. Gene and repeat annotations.

|  | **Col-0** | **An-1** | **C24** | **Cvi-0** | **Eri-1** | **Kyo** | **L*er*** | **Sha** |
| --- | --- | --- | --- | --- | --- | --- | --- | --- |
| **Protein-coding genes (Araport11)** | 27,445 | 27,342 | 27,214 | 27,098 | 27,285 | 27,574 | 27,376 | 27,293 |
| **miRNAs** | 298 | 316 | 301 | 297 | 301 | 313 | 296 | 301 |
| **snRNAs** | 75 | 76 | 78 | 76 | 77 | 77 | 78 | 75 |
| **snoRNAs** | 551 | 602 | 529 | 553 | 571 | 578 | 562 | 580 |
| **rRNAs** | 550 | 501 | 347 | 661 | 387 | 688 | 395 | 1,011 |
| **tRNAs** | 661 | 564 | 545 | 585 | 589 | 561 | 571 | 511 |
| **other RNAs** | 75 | 41 | 43 | 48 | 48 | 43 | 41 | 26 |
| **DNA transposons** | 10,278 | 10,095 | 10,452 | 10,769 | 10,834 | 11,265 | 10,689 | 10,702 |
| **retrotransposons** | 8,598 | 8,797 | 8,493 | 8,724 | 8,601 | 8,790 | 8,570 | 9,072 |
| **DNA transposon content** | 4.96% | 4.79% | 4.77% | 4.81% | 4.99% | 4.99% | 4.81% | 4.78% |
| **retrotransposon content** | 8.28% | 8.86% | 8.40% | 8.23% | 8.81% | 8.62% | 8.50% | 8.56% |
| **Complete repeat content** | 17.55% | 17.94% | 17.42% | 17.47% | 18.28% | 18.57% | 17.93% | 18.42% |

**Note**: The non-coding RNA and repeat annotation were predicted with Infernal and RepeatMasker, respectively.

#### Supplementary Table 10. Number and total length of rearranged regions.

| ***Number*** | |  |  |  |  |  |  |
| --- | --- | --- | --- | --- | --- | --- | --- |
| **Accession** | **INV** | **ITX** | **CTX** | **DUP-Loss** | **RefSp.** | **DUP-Gain** | **AccSp.** |
| An-1 | 33 | 364 | 365 | 2,181 | 5,551 | 2,107 | 5,805 |
| C24 | 37 | 473 | 508 | 2,423 | 6,305 | 2,207 | 6,455 |
| Cvi-0 | 37 | 566 | 626 | 2,546 | 7,478 | 2,604 | 7,705 |
| Eri-1 | 42 | 410 | 462 | 2,134 | 5,718 | 2,057 | 6,293 |
| Kyo | 33 | 368 | 430 | 2,026 | 5,681 | 2,277 | 6,516 |
| L*er* | 46 | 509 | 485 | 2,409 | 6,085 | 2,470 | 6,509 |
| Sha | 37 | 460 | 487 | 2,347 | 6,391 | 2,631 | 6,837 |
| ***Length*** | |  |  |  |  |  |  |
| An-1 | 1,824,867 | 985,071 | 862,756 | 3,861,135 | 5,320,292 | 3,377,683 | 5,171,246 |
| C24 | 1,742,898 | 1,367,338 | 1,311,211 | 4,562,135 | 5,929,351 | 3,301,993 | 5,900,176 |
| Cvi-0 | 1,485,583 | 1,657,606 | 1,273,284 | 4,417,754 | 6,345,126 | 4,074,912 | 6,294,286 |
| Eri-1 | 1,709,695 | 975,394 | 1,016,615 | 4,063,236 | 5,405,177 | 3,323,824 | 5,730,514 |
| Kyo | 1,559,482 | 988,729 | 1,147,826 | 3,917,699 | 5,056,775 | 3,790,715 | 5,511,508 |
| L*er* | 1,799,626 | 1,402,826 | 1,102,662 | 4,284,186 | 5,600,844 | 4,005,952 | 5,768,803 |
| Sha | 4,214,442 | 1,254,696 | 1,181,706 | 4,408,231 | 6,526,286 | 4,291,496 | 6,018,238 |

**Note:** INV: inversion; ITX: intra-chromosome translocation; CTX: inter-chromosome translocation; DUP-Loss: duplication loss in accession (i.e.: the reference Col-0 has more copies); DUP-Gain: duplication gain in accession (i.e.: the accession has more copies); RefSp: regions specific to the reference genome; AccSp: regions specific to the accession.

#### Supplementary Table 11. The percent of genomic rearrangements which reside in or overlap with pericentromeric regions

| **Genome** | **INV** | **ITX** | **CTX** | **DUP-Loss** | **DUP-Gain** | **Total** |
| --- | --- | --- | --- | --- | --- | --- |
| An-1 | 86.2% | 47.4% | 41.8% | 52.3% | 56.6% | 52.7% |
| C24 | 76.0% | 49.5% | 45.4% | 56.7% | 56.3% | 54.8% |
| Cvi-0 | 89.3% | 51.8% | 38.4% | 52.4% | 56.1% | 52.2% |
| Eri-1 | 85.0% | 48.9% | 45.6% | 55.3% | 55.5% | 53.7% |
| Kyo | 86.6% | 50.0% | 47.6% | 52.3% | 55.9% | 53.1% |
| L*er* | 82.5% | 59.0% | 48.1% | 59.6% | 62.2% | 59.4% |
| Sha | 61.5% | 52.7% | 41.4% | 54.8% | 61.3% | 56.1% |
| total | 45.3% | 51.6% | 43.8% | 54.8% | 57.8% | 54.6% |

**Note**: INV: inversion; ITX: intra-chromosome translocation; CTX: inter-chromosome translocation; DUP-Loss: duplication in reference but not in the accession genome; DUP-Gain: duplication in the accession genome but not in the reference genome.

#### Supplementary Table 12. Inversions larger than 50kb.

| **Accession** | **Chr** | **Start** | **End** | **Reference Length** | **Chr** | **Start** | **End** | **Query Length** |
| --- | --- | --- | --- | --- | --- | --- | --- | --- |
| An-1 | Chr1 | 14,501,429 | 15,074,096 | 572,667 | Chr1 | 14,867,422 | 15,215,011 | 347,589 |
| Sha | Chr1 | 17,660,763 | 17,879,638 | 218,875 | Chr1 | 17,552,872 | 17,764,136 | 211,264 |
| C24 | Chr1 | 23,188,572 | 23,504,543 | 315,971 | Chr1 | 22,126,192 | 22,444,978 | 318,786 |
| L*er* | Chr2 | 12,661,878 | 12,731,490 | 69,612 | Chr2 | 12,361,365 | 12,433,285 | 71,920 |
| Sha | Chr3 | 2,774,870 | 5,252,859 | 2,477,989 | Chr3 | 2,798,243 | 5,282,580 | 2,484,337 |
| L*er* | Chr3 | 8,295,153 | 8,465,270 | 170,117 | Chr3 | 8,351,497 | 8,521,688 | 170,191 |
| An-1 | Chr3 | 12,482,569 | 12,618,513 | 135,944 | Chr3 | 12,419,542 | 12,562,554 | 143,012 |
| An-1 | Chr3 | 13,602,813 | 13,921,313 | 318,500 | Chr3 | 13,323,011 | 13,379,922 | 56,911 |
| C24 | Chr3 | 13,602,813 | 13,924,434 | 321,621 | Chr3 | 14,225,825 | 14,285,314 | 59,489 |
| Eri-1 | Chr3 | 13,602,813 | 13,923,880 | 321,067 | Chr3 | 14,090,720 | 14,163,036 | 72,316 |
| Kyo | Chr3 | 13,602,813 | 13,921,313 | 318,500 | Chr3 | 14,288,380 | 14,342,210 | 53,830 |
| Sha | Chr3 | 14,336,990 | 14,554,838 | 217,848 | Chr3 | 13,546,260 | 13,785,045 | 238,785 |
| An-1 | Chr4 | 1,612,606 | 2,782,625 | 1,170,019 | Chr4 | 1,652,389 | 2,880,783 | 1,228,394 |
| Cvi-0 | Chr4 | 1,612,606 | 2,782,625 | 1,170,019 | Chr4 | 1,584,780 | 2,849,231 | 1,264,451 |
| Eri-1 | Chr4 | 1,612,606 | 2,782,625 | 1,170,019 | Chr4 | 1,690,889 | 2,870,416 | 1,179,527 |
| Kyo | Chr4 | 1,612,606 | 2,782,625 | 1,170,019 | Chr4 | 1,666,349 | 2,828,956 | 1,162,607 |
| L*er* | Chr4 | 1,612,606 | 2,782,625 | 1,170,019 | Chr4 | 1,746,529 | 2,898,561 | 1,152,032 |
| Sha | Chr4 | 1,612,606 | 2,782,625 | 1,170,019 | Chr4 | 1,655,631 | 2,860,573 | 1,204,942 |
| C24 | Chr4 | 1,771,815 | 2,782,625 | 1,010,810 | Chr4 | 1,628,501 | 2,773,489 | 1,144,988 |
| Sha | Chr4 | 4,356,088 | 4,459,766 | 103,678 | Chr4 | 4,583,599 | 4,658,107 | 74,508 |
| C24 | Chr4 | 4,572,269 | 4,692,550 | 120,281 | Chr4 | 5,054,623 | 5,231,295 | 176,672 |
| Eri-1 | Chr4 | 4,572,269 | 4,684,953 | 112,684 | Chr4 | 4,301,368 | 4,383,248 | 81,880 |
| Kyo | Chr4 | 4,573,080 | 4,692,550 | 119,470 | Chr4 | 4,533,759 | 4,689,570 | 155,811 |
| Sha | Chr5 | 11,730,380 | 11,918,318 | 187,938 | Chr5 | 11,786,872 | 11,986,397 | 199,525 |
| An-1 | Chr5 | 11,806,527 | 12,002,521 | 195,994 | Chr5 | 11,646,437 | 11,775,074 | 128,637 |
| An-1 | Chr5 | 12,437,159 | 13,049,679 | 612,520 | Chr5 | 12,282,399 | 12,909,890 | 627,491 |
| Eri-1 | Chr5 | 12,437,159 | 13,049,679 | 612,520 | Chr5 | 12,323,868 | 12,905,576 | 581,708 |
| Kyo | Chr5 | 12,437,159 | 12,989,685 | 552,526 | Chr5 | 12,188,346 | 12,650,586 | 462,240 |
| Sha | Chr5 | 12,437,159 | 12,989,685 | 552,526 | Chr5 | 12,846,278 | 13,225,872 | 379,594 |
| C24 | Chr5 | 12,477,370 | 13,049,679 | 572,309 | Chr5 | 11,887,516 | 12,455,714 | 568,198 |
| Cvi-0 | Chr5 | 12,539,575 | 13,049,679 | 510,104 | Chr5 | 12,289,995 | 12,635,489 | 345,494 |
| L*er* | Chr5 | 12,592,489 | 12,989,685 | 397,196 | Chr5 | 12,316,710 | 12,555,454 | 238,744 |
| L*er* | Chr5 | 13,050,974 | 13,284,321 | 233,347 | Chr5 | 12,555,453 | 12,810,352 | 254,899 |

#### Supplementary Table 13. Number and total length of local sequence variation in syntenic and rearranged regions.

| ***Number*** | | |  |  |  |  |  |
| --- | --- | --- | --- | --- | --- | --- | --- |
| **Accession** | **Region** | **SNP** | **S-Indel** | **L-Indel** | **HDR** | **CPG** | **CPL** |
| An-1 | SYN | 637,248 | 150,172 | 1,372 | 1,582 | 400 | 400 |
| An-1 | GR | 81,377 | 9,591 | 70 | 48 | 58 | 41 |
| C24 | SYN | 720,846 | 168,230 | 1,604 | 1,690 | 464 | 441 |
| C24 | GR | 103,207 | 12,332 | 95 | 62 | 60 | 56 |
| Cvi-0 | SYN | 892,125 | 209,003 | 1,851 | 2,282 | 522 | 541 |
| Cvi-0 | GR | 114,058 | 13,430 | 95 | 83 | 61 | 56 |
| Eri-1 | SYN | 682,249 | 159,143 | 1,438 | 1,582 | 441 | 439 |
| Eri-1 | GR | 84,278 | 9,696 | 73 | 53 | 55 | 50 |
| Kyo | SYN | 691,263 | 159,418 | 1,499 | 1,667 | 433 | 459 |
| Kyo | GR | 81,309 | 9,896 | 77 | 58 | 48 | 52 |
| L*er* | SYN | 693,354 | 161,088 | 1,439 | 1,687 | 447 | 467 |
| L*er* | GR | 102,048 | 11,952 | 83 | 56 | 52 | 53 |
| Sha | SYN | 727,954 | 170,344 | 1,533 | 1,793 | 465 | 491 |
| Sha | GR | 114,816 | 15,077 | 108 | 82 | 66 | 70 |
| ***Length*** | | |  |  |  |  |  |
| An-1 | SYN | 637,248 | 532,662 | 538,146 | 953,561 | 367,053 | 341,470 |
| An-1 | GR | 81,377 | 33,006 | 26,918 | 26,336 | 158,342 | 106,342 |
| C24 | SYN | 720,846 | 602,294 | 653,268 | 1,081,456 | 463,818 | 403,270 |
| C24 | GR | 103,207 | 41,370 | 20,938 | 44,956 | 288,835 | 150,235 |
| Cvi-0 | SYN | 892,125 | 742,298 | 723,616 | 1,556,755 | 551,227 | 492,673 |
| Cvi-0 | GR | 114,058 | 45,222 | 27,663 | 49,091 | 154,123 | 121,634 |
| Eri-1 | SYN | 682,249 | 570,014 | 566,821 | 972,935 | 393,342 | 469,857 |
| Eri-1 | GR | 84,278 | 32,141 | 27,734 | 27,278 | 139,160 | 209,105 |
| Kyo | SYN | 691,263 | 567,521 | 585,439 | 986,372 | 366,991 | 522,942 |
| Kyo | GR | 81,309 | 33,208 | 29,441 | 54,438 | 132,861 | 162,146 |
| L*er* | SYN | 693,354 | 571,606 | 558,989 | 946,557 | 489,357 | 481,803 |
| L*er* | GR | 102,048 | 40,617 | 22,175 | 38,796 | 120,753 | 224,427 |
| Sha | SYN | 727,954 | 609,894 | 606,149 | 1,162,736 | 475,067 | 533,052 |
| Sha | GR | 114,816 | 51,339 | 41,818 | 40,173 | 144,841 | 202,717 |

**Note:** SYN: syntenic regions; GR: rearranged regions including inversion, translocation and duplications; S-Indel: short indels; L-Indel: long indels; HDR: highly divergent regions; CPG: copy gain variation; CPL: copy-loss variation

#### Supplementary Table 14. Gene family contraction and expansion between Col-0 and each of the other seven genomes.

| **Acc.** | **NvN** | **1v2** | **2v1** | **1v3** | **3v1** | **2v3** | **3v2** | **Ref>Qry** | **Qry>Ref** | **ColSp** |
| --- | --- | --- | --- | --- | --- | --- | --- | --- | --- | --- |
| An-1 | 25,561 | 155 | 330 | 12 | 36 | 46 | 138 | 358 | 131 | 678 |
| C24 | 25,228 | 185 | 402 | 13 | 81 | 72 | 168 | 450 | 147 | 699 |
| Cvi-0 | 25,019 | 151 | 444 | 16 | 84 | 100 | 207 | 452 | 149 | 823 |
| Eri-1 | 25,379 | 175 | 378 | 17 | 63 | 62 | 120 | 397 | 154 | 704 |
| Kyo | 25,556 | 168 | 358 | 19 | 45 | 56 | 147 | 381 | 102 | 613 |
| L*er* | 25,399 | 164 | 388 | 14 | 84 | 72 | 144 | 381 | 135 | 665 |
| Sha | 25,184 | 163 | 394 | 14 | 78 | 74 | 159 | 448 | 139 | 792 |

**Note**: Gene copy number variation between Col-0 and each of other accessions based on gene family clustering. Acc.: accessions; NvN: both the Col-0 and the query genomes have the same number of genes in the respective gene families. 1v2, 2v1, 1v3, 3v1, 2v3, 3v2 indicate the corresponding number of Col-0 genes and accession genes in a gene family. “Ref>Qry” and “Qry>Ref” refer to the number of Col-0 genes in cases where either the reference or a divergent accession has more genes (>3) in a particular gene family. ColSp: genes present in the reference Col-0 genome but absent in the accession genome.

#### Supplementary Table 15. Non-reference genes.

| **Acc.** | **Novel** | **Novel2** | **AccSp** | ***A. lyrata* ortholog** | **RNA-seq support** |
| --- | --- | --- | --- | --- | --- |
| An-1 | 539 | 403 | 136 | 166 | 164 |
| C24 | 590 | 406 | 184 | 198 | 236 |
| Cvi-0 | 598 | 384 | 214 | 175 | 179 |
| Eri-1 | 537 | 416 | 121 | 171 | n/a |
| Kyo | 533 | 411 | 122 | 177 | n/a |
| L*er* | 555 | 420 | 135 | 172 | 270 |
| Sha | 551 | 413 | 138 | 173 | 146 |
| Total | 1,941 | 891 | 1,050 | 452 |  |

**Note**: The values in the “Total” row depict the numbers of gene families of the genes in the individual accessions (i.e. the non-redundant gene numbers). Novel2: genes shared by at least two accessions. AccSp: genes specific to an accession. n/a: no RNA-seq data available for these genomes.

#### Supplementary Table 16. List of HOT regions.

See attached Supplementary.Tables.v1.xlsx files.

#### Supplementary Table 17. R gene copy number differences in R gene clusters

| **ID** | **Chr** | **Start** | **End** | **An-1** | **C24** | **Col-0** | **Cvi-0** | **Eri-1** | **Kyo** | **L*er*** | **Sha** |
| --- | --- | --- | --- | --- | --- | --- | --- | --- | --- | --- | --- |
| RC001 | Chr1 | 4,140,216 | 4,147,939 | 2 | 2 | 2 | 2 | 2 | 1 | 1 | 2 |
| RC002 | Chr1 | 4,174,875 | 4,182,593 | 2 | 2 | 2 | 2 | 2 | 2 | 2 | 2 |
| RC003 | Chr1 | 6,052,936 | 6,060,851 | 3 | 3 | 3 | 3 | 3 | 3 | 3 | 3 |
| RC004 | Chr1 | 9,433,393 | 9,446,223 | 2 | 2 | 2 | 2 | 2 | 1 | 2 | 2 |
| RC005 | Chr1 | 21,167,565 | 21,1­85,495 | 6 | 3 | 3 | 5 | 3 | 2 | 3 | 3 |
| RC006 | Chr1 | 21,345,445 | 21,359,872 | 3 | 4 | 3 | 3 | 3 | 2 | 3 | 3 |
| RC007 | Chr1 | 21,420,244 | 21,428,078 | 2 | 0 | 2 | 2 | 1 | 1 | 2 | 0 |
| RC008 | Chr1 | 21,690,642 | 21,704,982 | 3 | 2 | 3 | 2 | 2 | 2 | 3 | 2 |
| RC009 | Chr1 | 21,746,354 | 21,833,018 | 5 | 1 | 5 | 1 | 2 | 1 | 4 | 2 |
| RC010 | Chr1 | 22,551,271 | 22,560,757 | 3 | 3 | 2 | 1 | 2 | 3 | 2 | 1 |
| RC011 | Chr1 | 22,607,462 | 22,616,373 | 1 | 1 | 2 | 1 | 1 | 1 | 1 | 1 |
| RC012 | Chr1 | 23,494,643 | 23,502,321 | 0 | 2 | 2 | 0 | 2 | 0 | 2 | 0 |
| RC013 | Chr1 | 23,641,770 | 23,655,453 | 3 | 3 | 3 | 3 | 3 | 3 | 3 | 3 |
| RC014 | Chr1 | 23,701,408 | 23,716,206 | 3 | 3 | 3 | 4 | 3 | 4 | 4 | 4 |
| RC015 | Chr1 | 27,409,196 | 27,445,997 | 8 | 9 | 11 | 8 | 13 | 6 | 11 | 9 |
| RC016 | Chr2 | 7,410,704 | 7,427,119 | 3 | 3 | 3 | 3 | 3 | 3 | 3 | 3 |
| RC017 | Chr3 | 1,105,650 | 1,112,437 | 2 | 2 | 2 | 2 | 2 | 2 | 2 | 2 |
| RC018 | Chr3 | 4,851,990 | 4,861,293 | 2 | 2 | 2 | 0 | 2 | 2 | 2 | 2 |
| RC019 | Chr3 | 16,044,662 | 16,096,477 | 1 | 2 | 2 | 0 | 2 | 2 | 0 | 4 |
| RC020 | Chr3 | 16,195,848 | 16,221,722 | 12 | 7 | 2 | 5 | 4 | 4 | 8 | 2 |
| RC021 | Chr3 | 17,206,183 | 17,215,755 | 3 | 2 | 2 | 2 | 3 | 1 | 2 | 1 |
| RC022 | Chr3 | 19,121,687 | 19,130,705 | 2 | 2 | 2 | 2 | 2 | 2 | 2 | 2 |
| RC023 | Chr4 | 5,940,186 | 5,973,113 | 3 | 0 | 3 | 0 | 3 | 3 | 0 | 3 |
| RC024 | Chr4 | 6,811,103 | 6,899,130 | 2 | 2 | 2 | 2 | 2 | 2 | 2 | 2 |
| RC025 | Chr4 | 7,196,991 | 7,209,692 | 2 | 2 | 2 | 2 | 2 | 2 | 2 | 2 |
| RC026 | Chr4 | 9,488,466 | 9,565,569 | 9 | 15 | 12 | 5 | 14 | 7 | 10 | 7 |
| RC027 | Chr4 | 10,439,984 | 10,446,300 | 3 | 3 | 2 | 2 | 3 | 4 | 2 | 2 |
| RC028 | Chr4 | 10,625,562 | 10,657,282 | 4 | 4 | 4 | 6 | 6 | 4 | 4 | 4 |
| RC029 | Chr4 | 10,795,869 | 10,800,792 | 3 | 3 | 3 | 0 | 3 | 3 | 3 | 3 |
| RC030 | Chr4 | 12,236,593 | 12,272,940 | 3 | 3 | 3 | 3 | 3 | 3 | 3 | 3 |
| RC031 | Chr4 | 13,620,977 | 13,637,119 | 1 | 1 | 2 | 1 | 1 | 1 | 1 | 1 |
| RC032 | Chr4 | 17,098,612 | 17,108,825 | 2 | 3 | 2 | 2 | 2 | 2 | 2 | 2 |
| RC033 | Chr5 | 5,908,834 | 5,951,750 | 3 | 3 | 3 | 1 | 3 | 3 | 4 | 3 |
| RC034 | Chr5 | 6,074,069 | 6,089,041 | 3 | 3 | 3 | 2 | 2 | 2 | 2 | 2 |
| RC035 | Chr5 | 15,320,375 | 15,331,528 | 3 | 3 | 3 | 3 | 3 | 3 | 3 | 3 |
| RC036 | Chr5 | 16,034,441 | 16,047,572 | 3 | 3 | 3 | 3 | 3 | 3 | 3 | 3 |
| RC037 | Chr5 | 16,612,659 | 16,620,824 | 2 | 1 | 2 | 1 | 2 | 2 | 2 | 2 |
| RC038 | Chr5 | 16,688,626 | 16,699,028 | 1 | 1 | 2 | 1 | 2 | 2 | 2 | 2 |
| RC039 | Chr5 | 17,560,179 | 17,569,148 | 2 | 2 | 2 | 1 | 1 | 1 | 2 | 1 |
| RC040 | Chr5 | 18,114,461 | 18,205,000 | 8 | 8 | 10 | 10 | 8 | 8 | 8 | 8 |
| RC041 | Chr5 | 18,283,663 | 18,332,920 | 11 | 8 | 7 | 5 | 9 | 11 | 9 | 11 |
| RC042 | Chr5 | 18,412,059 | 18,432,430 | 4 | 5 | 2 | 3 | 3 | 4 | 5 | 2 |
| RC043 | Chr5 | 18,758,699 | 18,769,090 | 2 | 2 | 2 | 2 | 2 | 2 | 2 | 2 |
| RC044 | Chr5 | 18,835,605 | 18,872,494 | 4 | 2 | 5 | 3 | 3 | 4 | 4 | 3 |
| RC045 | Chr5 | 19,185,698 | 19,195,559 | 2 | 2 | 3 | 2 | 3 | 3 | 2 | 2 |
| RC046 | Chr5 | 19,773,117 | 19,779,815 | 0 | 2 | 2 | 0 | 2 | 0 | 2 | 0 |
| RC047 | Chr5 | 26,712,944 | 26,721,377 | 3 | 3 | 3 | 3 | 3 | 3 | 3 | 3 |

### Supplementary Figures

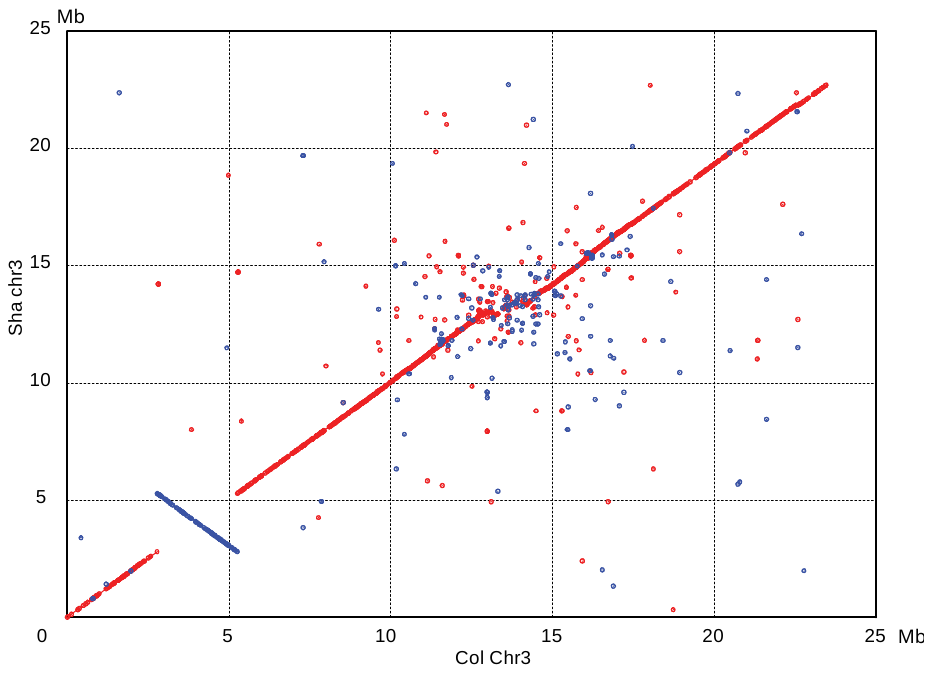

#### Supplementary Figure 1. A 2.48Mb inversion between the Col-0 reference and the Sha assembly.

The alignment was performed with nucmer with the parameter setting “-mum -c 40 -b 90 -l 100 -b 200”. The plot was drawn using MUMmerplot ^29^. Red: forward alignments, Blue: reverse alignments.

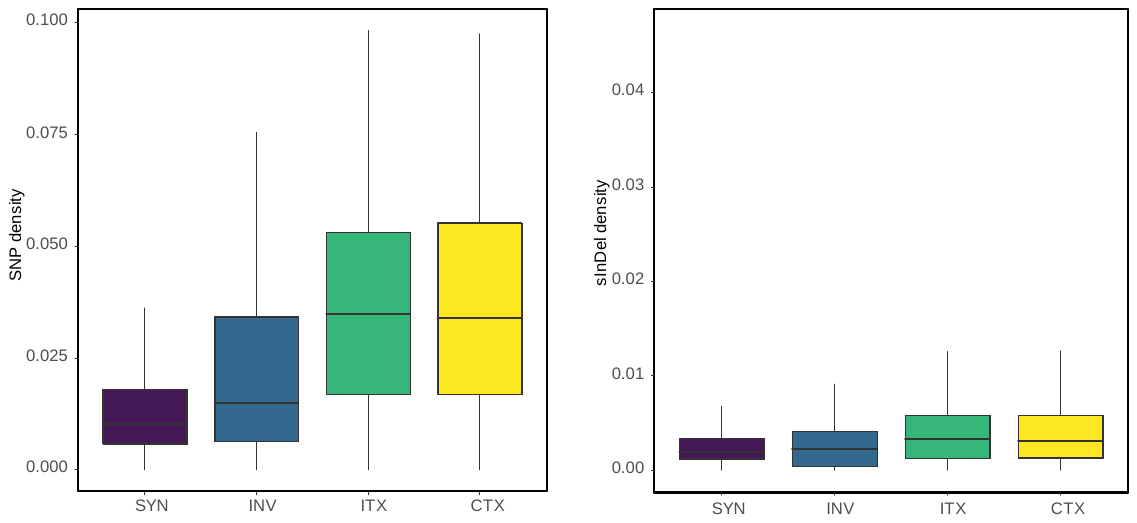

#### Supplementary Figure 2. Sequence variation (SNPs and small InDels) density per bp in syntenic and rearranged regions.

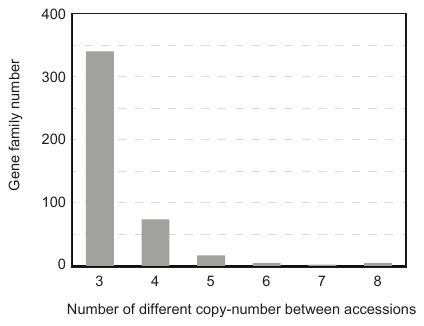

#### Supplementary Figure 3. Gene families with multiple different copy numbers across the eight accessions.

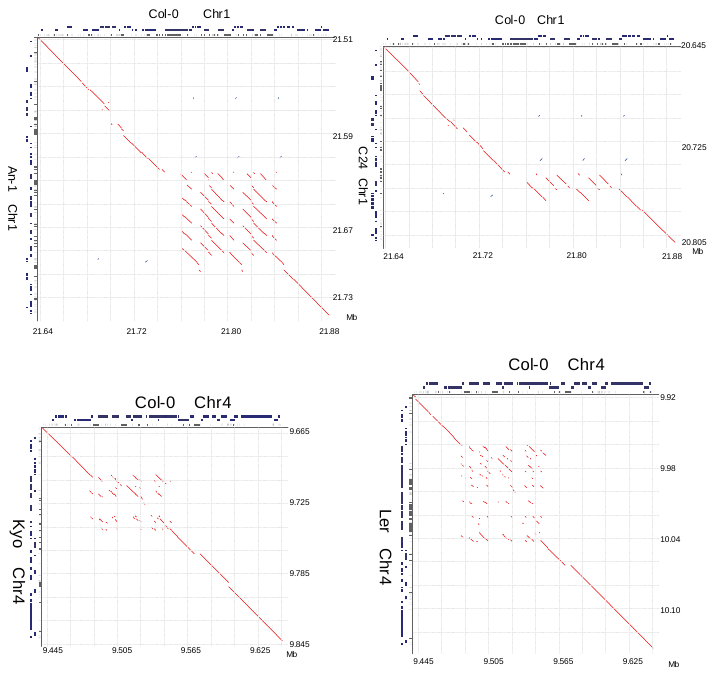

#### Supplementary Figure 4. Examples of sequence alignment dot plots of four hotspots of rearrangements. The three rows on top and the three columns on the right show the location of genes on the forward strand (top), on the reverse strand (middle) and the repeat regions (bottom). Red line: forward alignment, blue line: reverse alignment

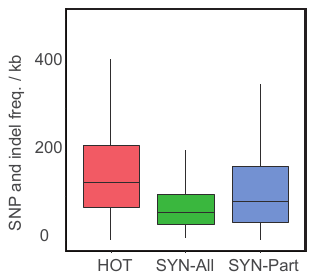

#### Supplementary Figure 5. Small variation frequency in HOT, SYN-All and SYN-Part regions.

The small sequence variation was identified based on the whole genome alignments between each of seven accessions and Col-0 reference genome. HOT: regions with Synteny Diversity values of larger than 0.5; SYN-All: regions which are syntenic in all pairwise comparisons (Synteny Diversity = 0); SYN-Part: partially syntenic regions, including all regions remaining after excluding SYN-All and HOT regions.

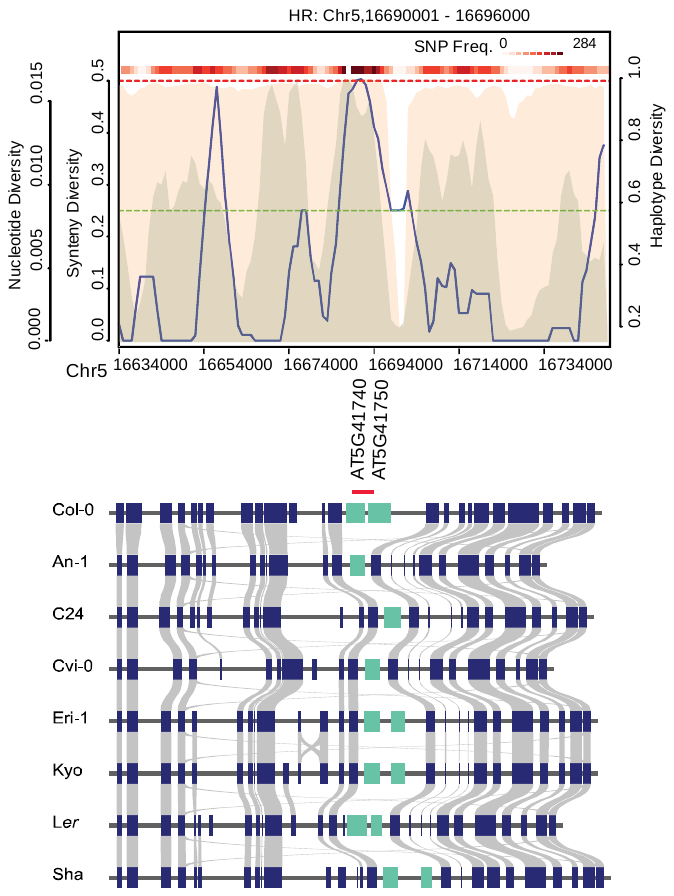

#### Supplementary Figure 6. Gene arrangements as well as Synteny, Nucleotide, and Haplotype Diversity in the *Dangerous Mix 1* (*DM1*) locus.

*DM1* contains two tandem duplicated genes AT5G41740 and AT5G41750 ^30^. One HOT region (shown by a red line above Col-0 genes) overlaps with *DM1*.

­

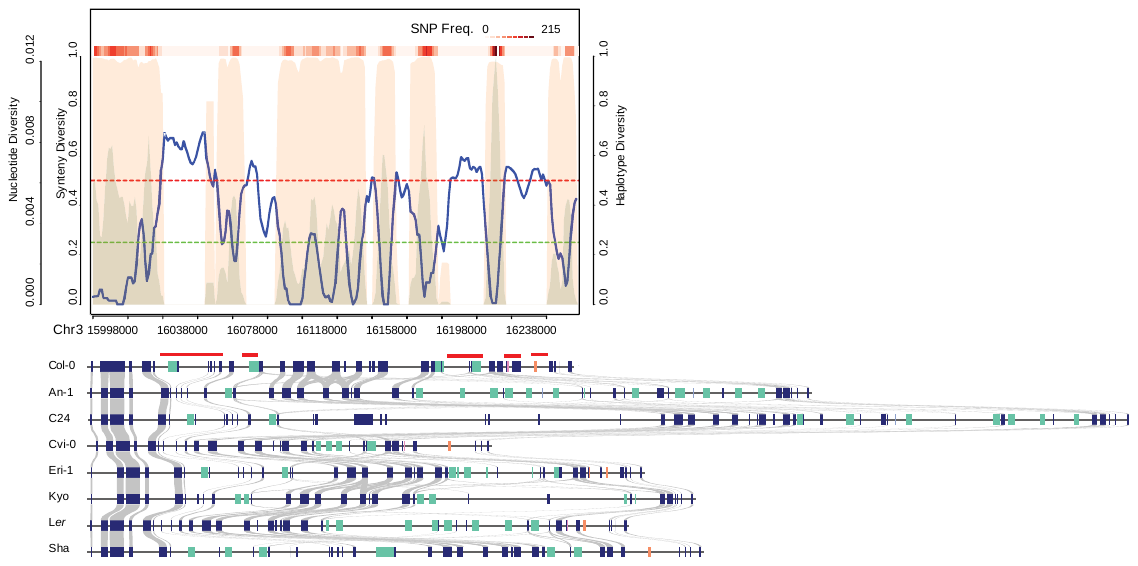

#### Supplementary Figure 7. Gene arrangements as well as Synteny, Nucleotide, and Haplotype Diversity in the *Dangerous Mix 2* (*DM2*) locus.

*DM2* was mapped to chromosome 3:16.15 - 16.30 Mb ^31^. Several HOT regions (shown by red lines above Col-0 genes) are in or around this locus.

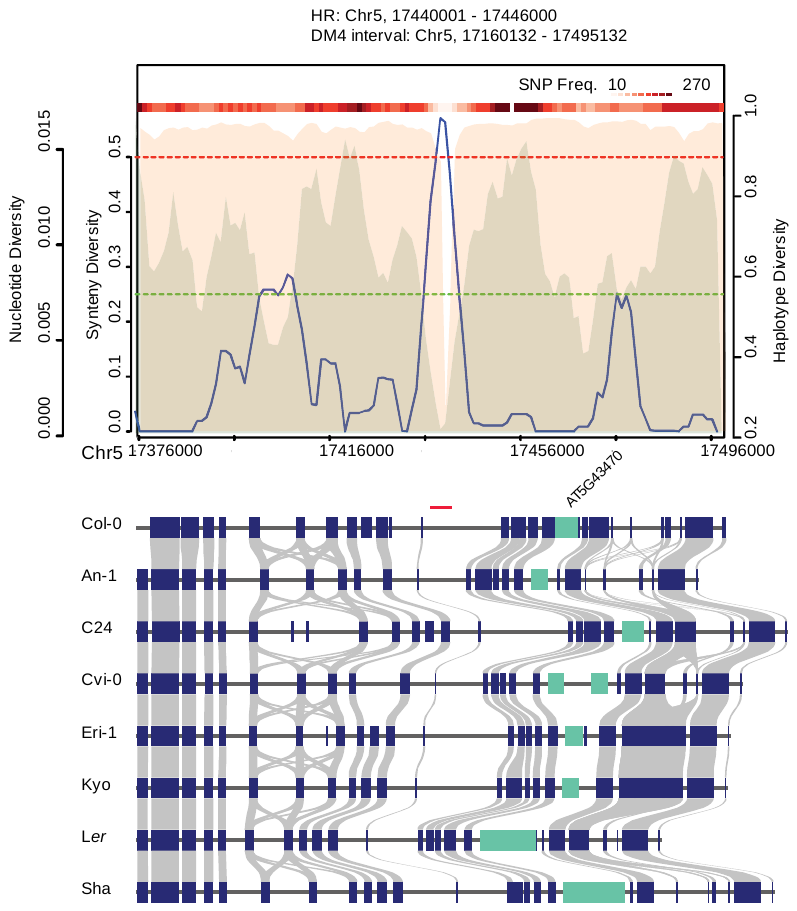

#### Supplementary Figure 8. Gene arrangements as well as Synteny, Nucleotide, and Haplotype Diversity in the *Dangerous Mix 4* (*DM4*) locus.

*DM4* locus was mapped to chromosome 5:17.16 - 17.50 Mb ^31^. One HOT region (shown by a red line above Col-0 genes) is in this locus.

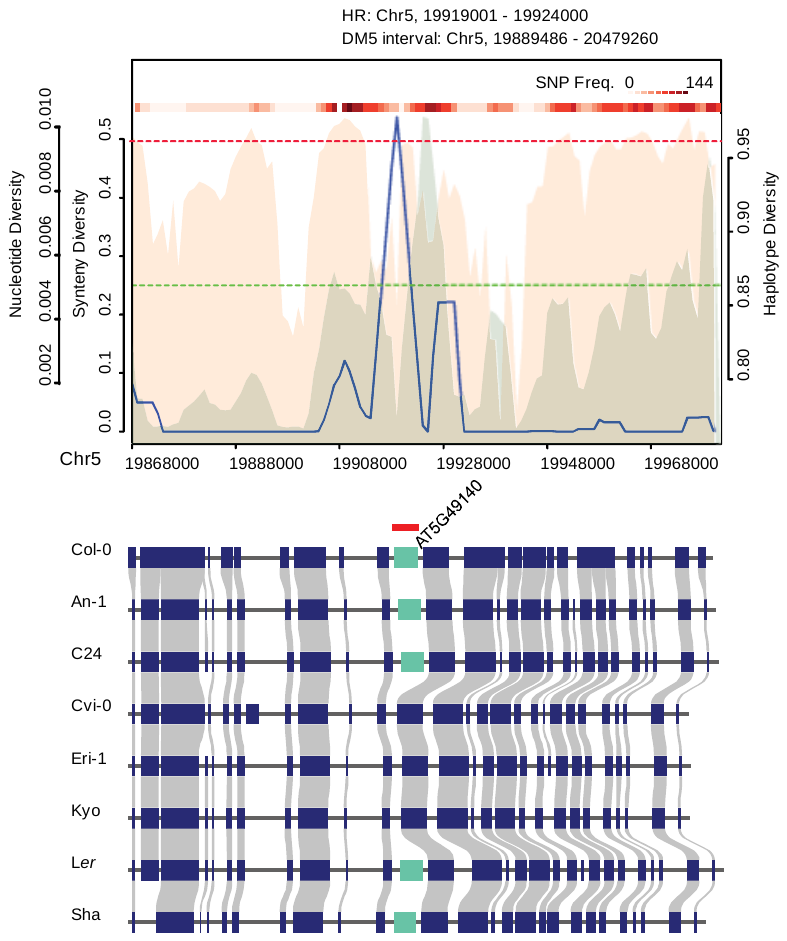

#### Supplementary Figure 9. Gene arrangements as well as Synteny, Nucleotide, and Haplotype Diversity in the *Dangerous Mix 5* (*DM5*) locus.

*DM5* locus was mapped to chromosome 5:19.89 - 20.48 Mb ^31^. One HOT region (shown by red line above Col-0 genes) is in this locus.

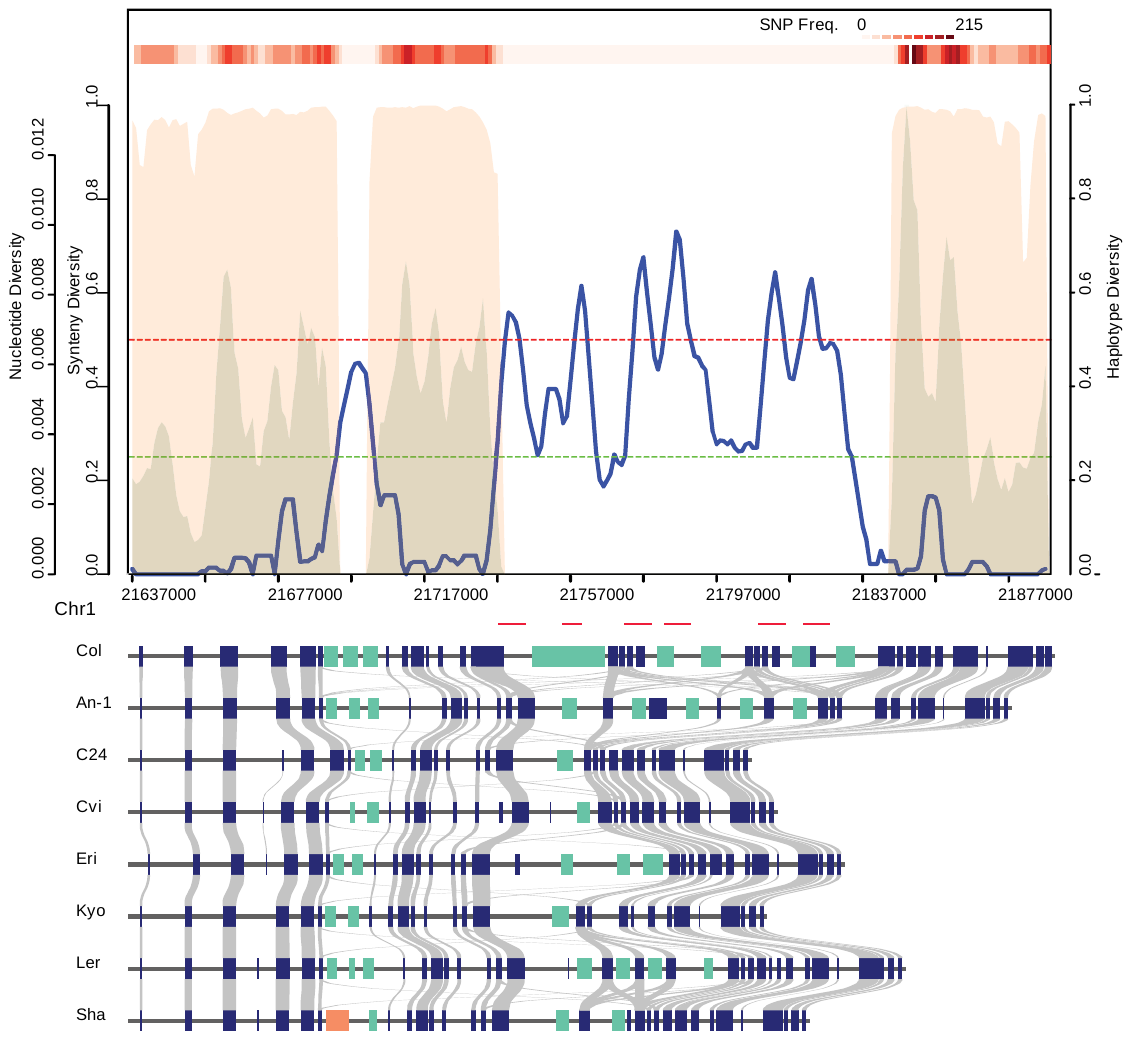

#### Supplementary Figure 10. Gene arrangements and synteny, nucleotide, haplotype diversity in dangerous mix (*DM*) locus *DM6.* The DM6 locus was mapped at chromosome 1 21.41 - 22.38 Mb ^31^. Several HOT regions overlapped with this interval.

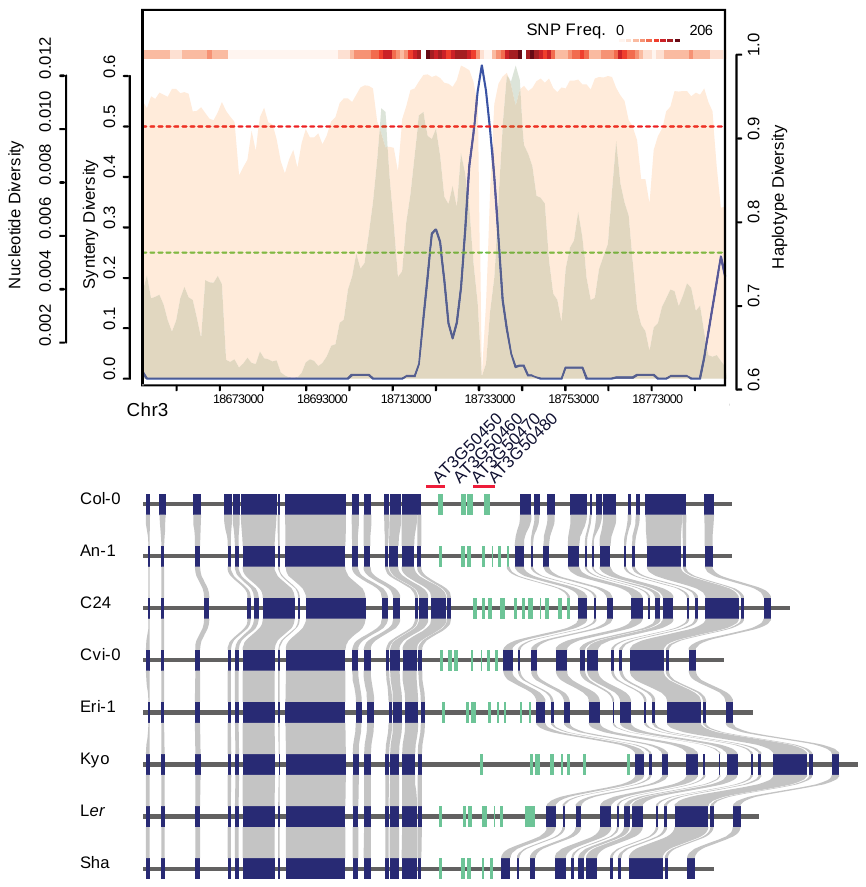

#### Supplementary Figure 11. Gene arrangements as well as Synteny, Nucleotide, and Haplotype Diversity in the *Dangerous Mix 7* (*DM7*) locus.

The *DM7* locus was mapped to two intervals on chromosome 3:18.53 – 18.99 Mb and 18.58-18.93 Mb ^31^. Two HOT regions (shown by red lines above Col-0 genes) overlapped with this interval and the R gene cluster *RPW8.1/RPW8.2*.

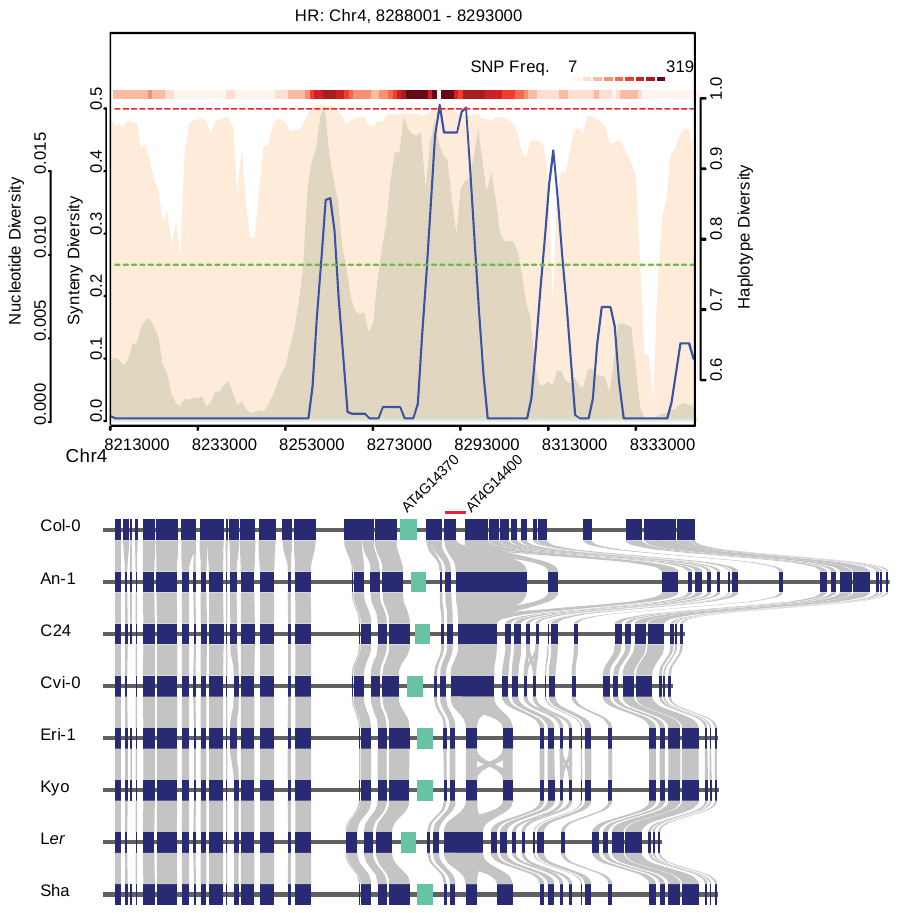

#### Supplementary Figure 12. Gene arrangements as well as Synteny, Nucleotide, and Haplotype Diversity in the *Dangerous Mix 9* (*DM9*) locus.

A small HOT region (shown by a red line above Col-0 genes) overlaps with the *DM9* *ACD6* allele (AT4G14400) ^32^.

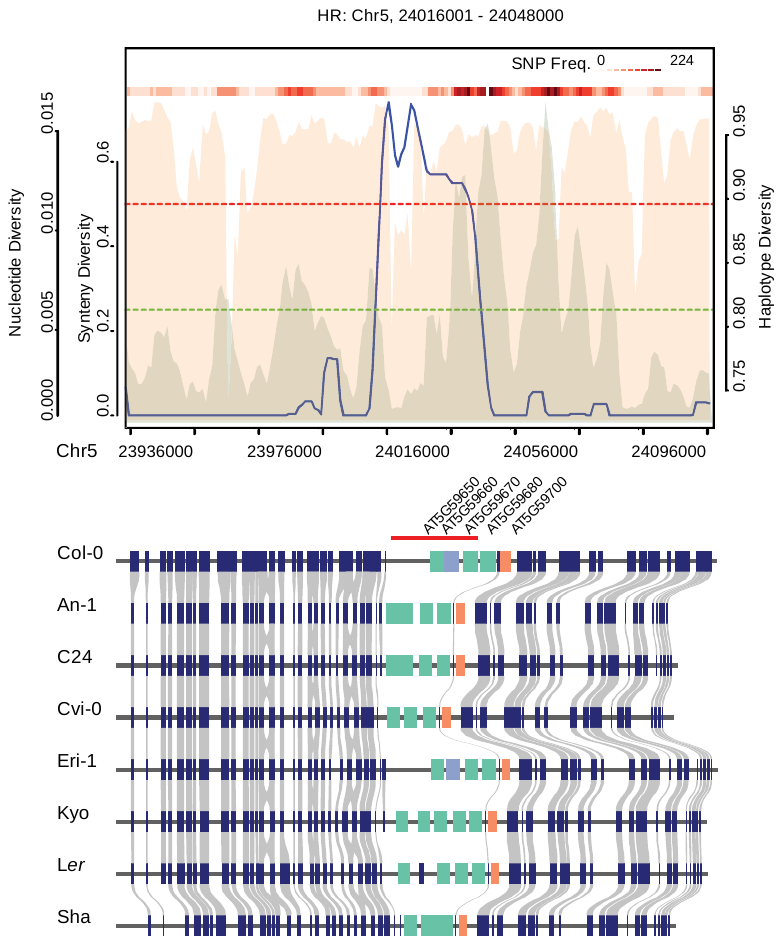

#### Supplementary Figure 13. Gene arrangements as well as Synteny, Nucleotide, and Haplotype Diversity in the single-locus genetic incompatibility locus of the OAK gene.

A large HOT region (shown by a red line above Col-0 genes) overlaps with the *OAK* gene including four receptor-like kinases (AT5G59650 to AT5G59680) ^33^.
